# Cancer cells have distinct electrical properties that predict a susceptibility to lipophilic anions; a new cancer drug paradigm

**DOI:** 10.1101/035113

**Authors:** Michael D. Forrest

## Abstract

I use the Nernst equation, parameterised with experimental data, to predict that cancer cells will accumulate more of a lipophilic anion than normal cells. This effect is correlated to charge number. Model cancer cells accumulate *100 more of an anion, *10^3^ more di-anion, *10^6^ more tri-anion, *10^8^ more tetra-anion and *10^10^ more penta-anion (>>1 billion times more). The trend endures, conveying even greater specificity, for higher charge numbers. This effect could be leveraged for cancer therapy. Wherein the lipophilic anion is a toxin that targets some vital cellular process, which normal and cancer cells may even share. It delivers a high, lethal dose to cancer cells but a low, safe dose to normal cells. This mathematical finding conveys the prospect of a broad, powerful new front against cancer.

## BACKGROUND

President Richard Nixon declared war on cancer over 40 years ago. However, today cancer is the leading cause of death worldwide. 7.6 million deaths in 2008; projected to rise to 13.1 million deaths per year by 2030 [1]. It is estimated that a quarter of adult males, and a fifth of female adults, will die from cancer in the USA [2]. These fatalities deliver substantial economic and immeasurable personal cost.

I propose a new class of anti-cancer medicines. Lipophilic anions: negatively-charged molecules that can passage through lipid membranes. I use a biophysical model to show that cancer cells will accumulate and retain more of these molecules, in their cytoplasm and mitochondrial intermembrane space (IMS), than normal cells. So, a poison from this class will be directed to cancer cells at a higher dose than normal cells, which will convey a therapeutic window. The model predicts that the accumulation by cancer cells is so much greater than for normal cells that even molecules/processes that normal cells rely on also are safe targets. So the potential of this new paradigm is immensely rich and broad, given the huge number of different molecular targets in the cytoplasm and IMS.

My accumulation model is a simple and powerful recasting of the Nernst equation [3], a cornerstone of biophysics, parameterised to experimental data. In the model, cancer and normal cells accumulate charged lipophilic molecules differently because they have different voltages across their plasma (Ψ_PM_) and inner mitochondrial (Ψ_IM_) membranes. As compared to normal cells, cancer cells have a depolarised Ψ_PM_ [4-10] and a hyperpolarised Ψ_IM_ [11-23]. Whether they have a different voltage across their outer mitochondrial membrane (Ψ_OM_) is less clear, with the relevant experiments yet to be done. The ramifications of different Ψ_OM_ values are explored in the model, but no tested Ψ_OM_ settings change model behaviour sufficiently to change the conclusions drawn.

There is a history of charged lipophilic molecules in cancer research. But cations rather than anions: delocalised lipophilic cations (DLCs) targeted to the mitochondrial matrix. A number of different DLCs have been shown to accumulate in, and selectively kill, cancer cells *in vitro* and *in vivo* [11-16, 20, 24-40]. However, no DLC has yet been successful in clinical trials. MKT-077 failed because of renal toxicity [41-42]. In phase I trials, rhodamine 123 couldn’t kill cancer cells, to the stringency of statistical significance, at the maximally tolerated dose [43]. These failures seem to have blunted an approach that once generated such excitement, promise and hope. To date, it hasn’t been explained why DLCs could be so successful in some cases (e.g. xenograft mouse models) and not others (e.g. clinical trials). My model delivers answers. It gives a mechanistic account of why some cancer cell lines will selectively accumulate more of a DLC than normal cells, and not others. A hyperpolarised Ψ_IM_ favours, and a depolarised Ψ_PM_ disfavours, DLC accumulation. For some cancers, their stereotypical depolarisation in Ψ_PM_ equals their stereotypical hyperpolarisation in Ψ_IM_. This cancels out any selective accumulation of DLCs as compared to normal cells. For other cancers, there is a small discrepancy between the magnitude of their stereotypical depolarisation in Ψ_PM_ and hyperpolarisation in Ψ_IM_. This conveys a margin for selective accumulation of DLCs over and above normal cells. By understanding why the DLC approach won’t work in every case, I hope it will reignite this tact for some amenable cancers. However, where the model explains why DLCs have a delimited therapeutic potential, it shows that lipophilic anions will have a universal applicability. Whereas the depolarisation in Ψ_PM_ and hyperpolarisation in Ψ_IM_ are subtractive for cations, they are additive for anions. And where the selective accumulation for DLCs is small at best, the selective accumulation of lipohilic anions is substantial at worst. So, I propose that lipophilic anions should be our focus rather than lipophilic cations; delocalised lipophilic anions (DLAs) as a new therapeutic paradigm.

### Cancer cells have a depolarised Ψ_PM_ compared to normal cells

Ψ_PM_ can change during the different phases of the cell proliferation cycle [4]. For example, Ψ_PM_ hyperpolarises in some cancer cells as they enter the S phase. However, at all phases, the Ψ_PM_ of cancer cells is consistently depolarised to that of normal, differentiated cells [4]. For example, MCF-7 breast cancer cells have a Ψ_PM_ value of −9 mV (-30 mV during S phase) and normal, differentiated breast cells have a Ψ_PM_ value of −40 to −58 mV. Ψ_PM_ depolarisation seems to be required for cancer proliferation (causative rather than merely correlative [4-5, 7]). For proliferating Chinese hamster ovary (CHO) cells, Ψ_PM_ = −10 mV, and hyperpolarising the cells to −45 or −75 mV (values more typical to normal differentiated cells) halts their proliferation [4]. So, strategies or drugs to hyperpolarise Ψ_PM_ may be effective against cancer. Indeed, a hyperpolarisation of Ψ_PM_ reduces tumour formation *in vivo* [5]. Ψ_PM_ values for cancer cells are depolarised to −40 mV; Ψ_PM_ values for normal, differentiated somatic cells are hyperpolarised to −40 mV (Figure 1 in [4]). Exact values vary between cell types. A fair, representative Ψ_PM_ value for a cancer cell = −10 mV; a fair, representative Ψ_PM_ value for a normal adult cell = −70 mV [4]. I will carry these values into my model.

**Figure 1.**
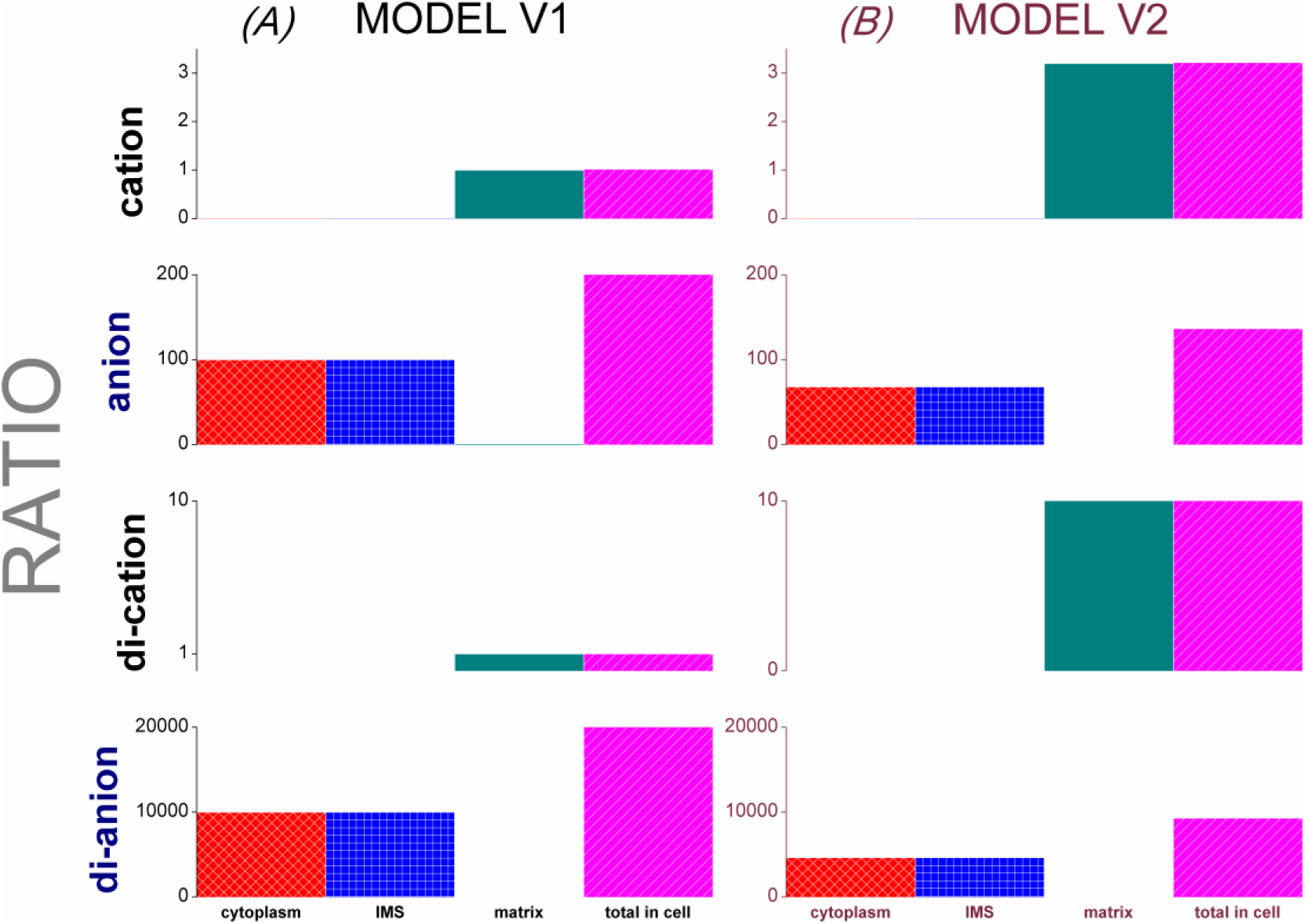
The biophysical model predicts that cancer cells accumulate more of a lipophilic anion than normal cells. (*A*) x-axis presents intracellular compartments, y-axis shows *Ratio* (differential in accumulation between cancer and normal cells). Cancer (Model V1) and normal cells have equal DLC accumulation (cation, di-cation: *Ratio* = 1) but cancer has a markedly greater anion/di-anion accumulation. (*B*) Cancer (Model V2) accumulates slightly more DLC (cation, di-cation) and much more anion/di-anion than a normal cell.

Note that there is a divergence and contradiction in the literature, with many hundreds of studies (e.g. the model of [39]) propagating an earlier report that cancer cells have a hyperpolarised, and not a depolarised, Ψ_PM_ compared to normal cells [16]. This report increased extracellular [K^+^] to depolarise the cell and identify the importance of Ψ_PM_. However, it doesn’t compare a cancer cell with its normal, untransformed equivalent. It compares a MCF-7 human breast cancer cell with a normal CV-1 African green monkey kidney epithelial cell line [16]. So, it doesn’t control for diversity in Ψ_PM_ between different cell types (and species). Nor for different cell types having a different depolarisation response to increased extracellular [K^+^], because of different ion channel complements. For example, different cell types can differ in their expression levels of pannexin (Cl^-^ channel), which has a greater open probability at hyperpolarised values of Ψ_PM_, in response to increased extracellular [K^+^] [44]. Studies that are more controlled, and that compare transformed and untransformed cells of the same origin, observe cancer cells to have a more depolarised Ψ_PM_ than normal cells [6]. In my modelling, I follow this conclusion, and it aligns with other studies, which also show Ψ_PM_ is depolarised in cancer cells e.g. [4-5, 7].

### Cancer cells have a hyperpolarised Ψ_IM_ compared to normal cells

In a normal cell, Ψ_IM_ flickers between −108 and −159 mV: with a mean value of −139 mV [45-46]. Cancer cells have a more hyperpolarised Ψ_IM_ than normal cells [11-23]. The more invasive and dangerous the cancer, the more hyperpolarised its Ψ_IM_ is observed to be [17-19]. The hyperpolarisation of Ψ_IM_ can be >50% greater in cancer cells than normal cells [17] e.g. Ψ_IM_ = ~-210 mV in Neu4145 cancer cells [21]. The Ψ_IM_ hyperpolarisation in cancer cells can even be double that of normal cells [22]. In a prior paper, I provided a quantitative, biophysical explanation of why Ψ_IM_ is more hyperpolarized in cancer cells [23].

### Cancer and normal cells may or may not have a different Ψ_OM_: data is limited and theory divergent

Experiments with pH sensitive probes, in cancer cells, show that the pH in the mitochondrial intermembrane space (IMS) is lower (6.9) than in the cytoplasm (7.6) [47]. If protons can’t move freely across the mitochondrial outer membrane (OM), this pH differential indicates Ψ_OM_ = +43 mV (Nernst equation, *T* = 310 K). However, protons can probably – I assume – flow through the OM via the voltage-dependent anion channel (VDAC). In which case, there must be a potential difference maintaining this pH difference, indicating Ψ_OM_ = −43 mV. On the other hand, the protons may not actually be free to move because of a boundary layer effect. [48] propose, on the basis of mitochondrial cristae anatomy/folding and thermodynamics [49], a boundary layer (~1 nm thick) on the IMS face of the mitochondrial inner membrane (IM) where the proton concentration is higher than the bulk (IMS and cytoplasm) concentration. If the protons are not free to leave the boundary layer, they certainly aren’t free to cross the OM. So, this would then indicate Ψ_OM_ = +43 mV again. Although to be accurate, here the potential is not across the OM but across the border of the boundary layer. But for our purposes here the exact location of Ψ is less important, than its value, which dictates how charged lipophilic molecules will move across it. This value may be an underestimate. It has been calculated that the pH in the boundary layer must be 2 pH points lower than the mitochondrial matrix, in order for the pmf to be large enough to produce sufficient ATP to support life [48]. The pH is ~8 in the mitochondrial matrix (7.8 [47], 8 [50]) and, by this argument, ~6 in the boundary layer. So, this calculation suggests the pH probe experiment [47] underestimates the pH difference between the IMS and cytoplasm. Indeed, we don’t know if this pH probe, attached to the C-terminus of an integral membrane protein in the IM, is sensing the pH in the boundary layer (which is still only a postulate) or at another point in the IMS. With this revised, estimated pH differential we attain Ψ_OM_ = +123 mV. Alternative theoretical studies, considering various different factors and in publication date order, have estimated Ψ_OM_ to be −5 mV [51], −60 to +60 mV [52], +12 mV [53], +30 mV at a maximum and −15 mV likely [54]. The latter is late enough to discuss the pH data of [47], but doesn’t frame it and consider it as I do here. This pH data is actually from a cancer cell line [47], we don’t have any comparative data from a normal, differentiated cell. Ψ_OM_ has physiological implication because VDAC, in the OM, is voltage-gated [55]. It has a bell-shaped conductance profile. It’s fully open when Ψ_OM_ is in the range: −40 to +40 mV, but partially closed at more positive/negative potentials than this. Its pore radius reduces from [1.2-1.5] to [0.85-0.95] nm [55] and it reverses its moderate selectivity, becoming more selective for cations than anions. In this state, ATP^4-^ passage is low [55].

My own view is that in addition to, or in place of, a potential difference across the OM: there may be a potential difference across the IMS facing border of a proton boundary layer (~1 nm thick) which resides along the IMS face of the IM. I propose that the potential difference due to a pH difference (= +43 to +123 mv) resides here, rather than at the OM, because at the OM this potential is sufficient to close VDAC, which would then pose the question of how ATP, ADP, NADH etc. passage the OM; for normal cells, and for cancer cells also. I think this is an important reanalysis and solves a paradox. However, in this paper I conform and speak only of Ψ_OM_. The exact location of this Ψ isn’t important to my model or study. Nor its value really: just whether it differs between normal and cancer cells. Incidentally, I envisage that this Ψ will “charge screen” the negative mitochondrial matrix and dampen the attraction of cations to it, but it will assist the localisation of anions to the IMS. This is another reason why I propose lipophilic anions rather than cations. This screening effect might be complex as protons may be distributed unequally in the IMS, preferentially located in cristae folds [49]. In which, [H^+^] is higher than bulk IMS. So, there will be areas of higher and lower screening.

In summary, we just don’t know what Ψ_OM_ is in normal and cancer cells, and if they differ at this parameter. I think an experiment in which the pH probe is attached to an integral protein of the OM rather than the IM (facing the IMS) would be extremely informative. However, for now, there is no meaningful consensus. So, with this uncertainty, my biophysical model uses Ψ_OM_ = 0 mV for both normal and cancer cells. However, I do explore the effect of different Ψ_OM_ values, and varied Ψ_OM_ differentials between normal and cancer cells, to investigate changes in model behaviour and if they can alter conclusions drawn. No Ψ_OM_ values tested undermine the findings of the paper. Indeed, both a positive or negative differential in Ψ_OM_ (between normal and cancer cells) can actually reinforce my findings. They make the selective accumulation of lipophilic anions by cancer cells reach even greater and even more stratospheric multiples. Some have suggested that Ψ_OM_ is more depolarised in cancer cells [56-57] in a theory that I summarise in the *Appendix*.

### A new drug design rule: the more anionic a drug candidate, the greater its probable selectivity for cancer cells

I use the model to suggest that anionic character should be a new selection criterion for cancer drug design. The more anionic, the better. This stipulation can be applied alone – or combined with Lipinski's rule of five [58], which predicts “drug-likeness” (oral bioavailability) – to generate a new drug design rule for cancer specific drugs. This can guide novel drug screens e.g. systematically testing drugs that conform to the rule of five (or some variant of it) in turn: from the most to the least anionic. This should find some of the best drugs early. Indeed, I have conducted such a screen myself. Excitingly, there are some very anionic compounds that conform to the rule of five. For example, anions with −6 charge e.g. C_14_H_22_O_10_P_2_^-6^ [59]. My model predicts that cancer cells will accumulate these >1 trillion times (!) more than normal cells. Some of the molecules I have found in my screens have had some bioactivity established, although not necessarily cancer-specific, by prior studies. But again, to repeat from earlier, the differential in accumulation is so great that targets which normal and cancer cells share are valid therapeutic targets for drugs of this class. For example, C_28_H_19_NO_15_S_3_^-4^ [60] disrupts histone modification [61]. This is an important process in transcriptional control and DNA packaging. My model predicts that this chemical, and its disruption, will be targeted to cancer cells 100 million times more than normal cells. Or there is C_28_H_17_N_5_O_14_S_4_^-4^ [62], which inhibits human Flap Endonuclease 1 ([63]; FEN1; very important in genomic stability). Again, my model predicts that this poison will be targeted to cancer cells 100 million times more than normal cells.

## MATERIALS AND METHODS

### The Model

Equations 1-3 use the Nernst equation [3] to present the lipophilic cation/anion concentration accumulated in the mitochondrial matrix (Eq. 1), mitochondrial intermembrane space (IMS) (Eq. 2) and cytoplasm (Eq. 3) of a cell as a function of the extracellular cation/anion concentration, [extracellular], which is arbitrarily set to 1. All equations were solved in R [65].

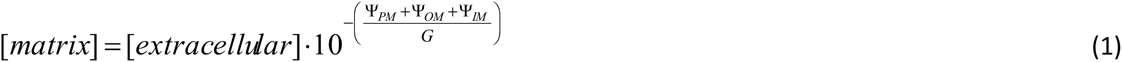

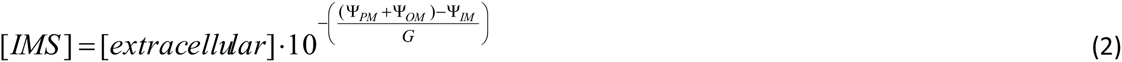

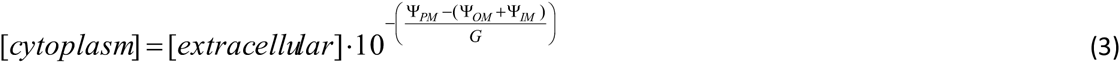

*G*=1000*2.3026*(RT/zF); where 1000 is a dimensionality factor to work in mV, *R* is the gas constant (8.31 *J* · *mol* ^1^), *T* is temperature (300 K), *z* is the charge of the species and *F* is the Faraday constant (96,400 *C* · *mol*^-1^). *G* = 60 for a cation, G = −60 for an anion. When cationic charge = −2, −3, −4, −5, −6, −7, −8, −9, −10, −20: *G* = 30, 20, 15, 12, 10, 8.5, 7.4, 6.6, 6, 3 respectively. Ψ_PM_, Ψ_OM_, Ψ_IM_ are the membrane potentials of the plasma, outer and inner mitochondrial membranes respectively. Their values can differ between cancer and normal cells. Thence, they accumulate lipophilic cations/anions differently. In my modelling:

Normal cell: Ψ_PM_ = −70 mV, Ψ_OM_ = 0 mV, Ψ_IM_ = −140 mV Cancer cell (Model V1): Ψ_PM_ = −10 mV, Ψ_OM_ = 0 mV, Ψ_IM_ = −200 mV Cancer cell (Model V2): Ψ_PM_ = −30 mV, Ψ_OM_ = 0 mV, Ψ_IM_ = −210 mV.

So, note that two different cancer model settings are trialled. Both have parameters within the experimentally observed range [4-10, 11-23]. So are equally valid. They reflect the delimited diversity in Ψ_PM_ and Ψ_IM_ between different cancers. Cancers, as compared to normal cells, consistently have a depolarised Ψ_PM_ [4-10] and hyperpolarised Ψ_IM_ [11-23] but the actual values involved within this trend can vary among different cancers. We capture some of this diversity by having two alternative cancer models, which differ in the degree of their Ψ_PM_ depolarisation and Ψ_IM_ hyperpolarisation: Models V1 and V2. The parameter values are chosen to give a reasonable account of the diversity.

Equations 1-3 were solved for a normal and a cancer cell, for different molecular species (cations and anions of different charge numbers: 1-10). Then, for each intracellular compartment (mitochondrial matrix, IMS, cytoplasm), the ratio of [concentration found in cancer cell vs. concentration found in normal cell] was calculated for these different species. This was done for both Model V1 and V2 as the cancer cell. It was also done for other more hypothetical normal and cancer cells, with different Ψ_PM_, Ψ_OM_, Ψ_IM_ values and less foundation in experiments, to explore parameter space. The *Ratio:* [accumulation in cancer cell, *x*]:[accumulation in normal cell, *y*] was calculated by:

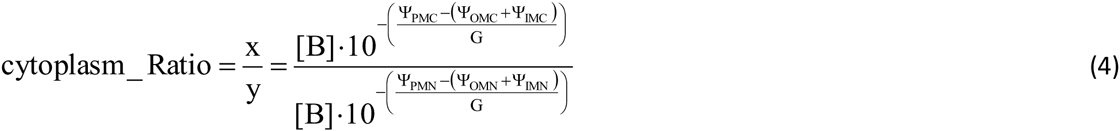

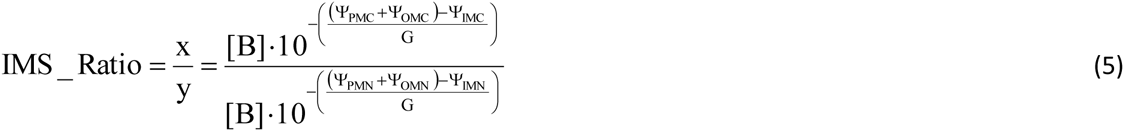

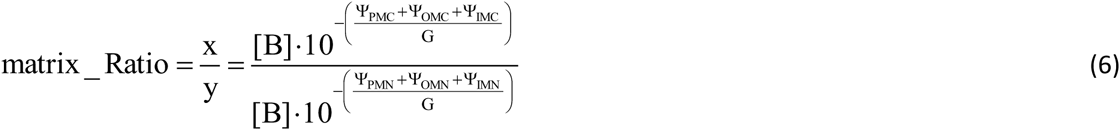

Where Ψ_PMC_, Ψ_OMC_, Ψ_IMC_ are cancer cell model values (Model V1 or V2), and Ψ_PMN_, Ψ_OMN_, Ψ_IMN_ are normal cell model values, for the plasma, outer mitochondrial and inner mitochondrial membrane potentials respectively. *G* is positive for a cationic species, negative for an anionic species.

Because of uncertainty about Ψ_OM_ values in normal and cancer cells (refer *Background*), Ψ_OM_ = 0 mV for all models at default. However, alternative Ψ_OM_ values are tested and reported to explore the parameter space. It is a differential in Ψ_OM_, between normal and cancer cells, that is salient and different (differential, Ψ_DIFF_) = (cancer Ψ_OM_) − (normal Ψ_OM_) values are trialled; across positive and negative values of this Ψ_OM_ differential.

## RESULTS

### Cancer cells accumulate lipophilic anions more than normal cells

Figure 1 presents model output; *A* shows that cancer (Model V1) and normal cells have equal DLC accumulation, both for cations and di-cations. By contrast, it shows that cancer cells (Model V1) accumulate *100 and *10,000 more lipophilic anions and di-anions respectively than normal cells; in both their cytoplasm and mitochondrial intermembrane space (IMS). Figure 1*B* shows an alternative cancer cell (Model V2) to have a slight accumulation of DLCs: *3 for cation, *10 for di-cation. By contrast, it shows that this cancer cell (Model V2) accumulates *68 and *4,642 more lipophilic anions and di-anions respectively than normal cells; in both its cytoplasm and mitochondrial intermembrane space (IMS). These computational results suggest that lipophilic anions are disproportionally targeted to cancer cells, better than DLCs.

How does the differential in anion/cation accumulation, between cancer and normal cells, scale with charge number?

Figure 1 only considered single and double charged species. Figure 2 shows the differential in cancer/normal cell accumulation of anions/cations, which differ in charge number (1-10). For cations, accumulation in the mitochondrial matrix is presented. For anions, accumulation in the IMS/cytoplasm (they are equal, refer Figure 1) is presented. It is clear again that anionic species are disproportionally accumulated by cancer cells, to a larger extent than cationic species. Indeed, Model V1 has no greater accumulation of any DLC, no matter its charge, than a normal cell (*Ratio* =1). Model V2 does show disproportional accumulation of DLCs. Models V1 and V2 show that the more anionic a molecule, the greater its accumulation by cancer cells as compared to normal cells. As mentioned in the *Introduction*, there are some very anionic molecules (-6) that still confirm to Lipinski’s rule of five for orally available drugs. Figure 2 shows that these (-6) will be accumulated in cancer cells >1 trillion times more than in normal cells. In the *Discussion* section, methodologies that can make use of very anionic molecules, significantly more negative than −6, are presented e.g. boron neutron capture therapy. Figure 2 shows that if a lipophilic molecule has 10 negative charges, it will be accumulated in cancer cells >>1 quintillion more times than normal cells.

**Figure 2.**
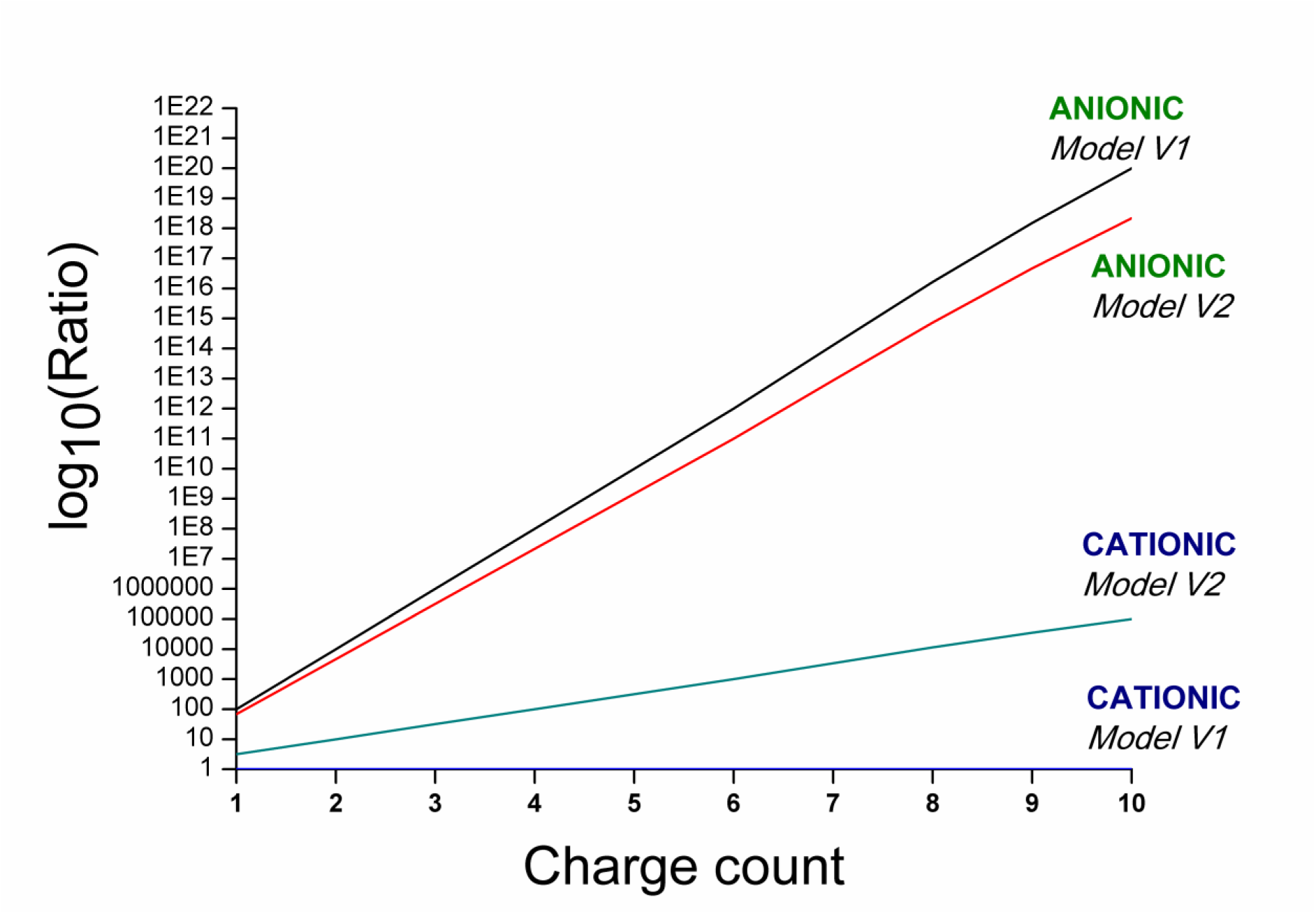
The differential in anion/cation accumulation, between cancer and normal cells (*Ratio*), scales with charge number. x-axis presents charge number, y-axis presents log_10_ of *Ratio*: the differential in accumulation between cancer and normal cells.

### Exploring some model parameter space

Figure 3; *A* shows the *Ratio* of cation accumulation in the mitochondrial matrix of a cancer cell, to that in a normal cell, as cancer Ψ_PM_ and Ψ_IM_ varies (Ψ_OM_ = 0 mV). *B* shows the *Ratio* of anion accumulation in the mitochondrial IMS (or cytoplasm) of a cancer cell, to that in a normal cell, as cancer Ψ_PM_ and Ψ_IM_ varies (Ψ_OM_ = 0 mV). The normal cell Ψ_PM_, Ψ_OM_, Ψ_IM_ parameters are as detailed in *Methods*.

**Figure 3.**
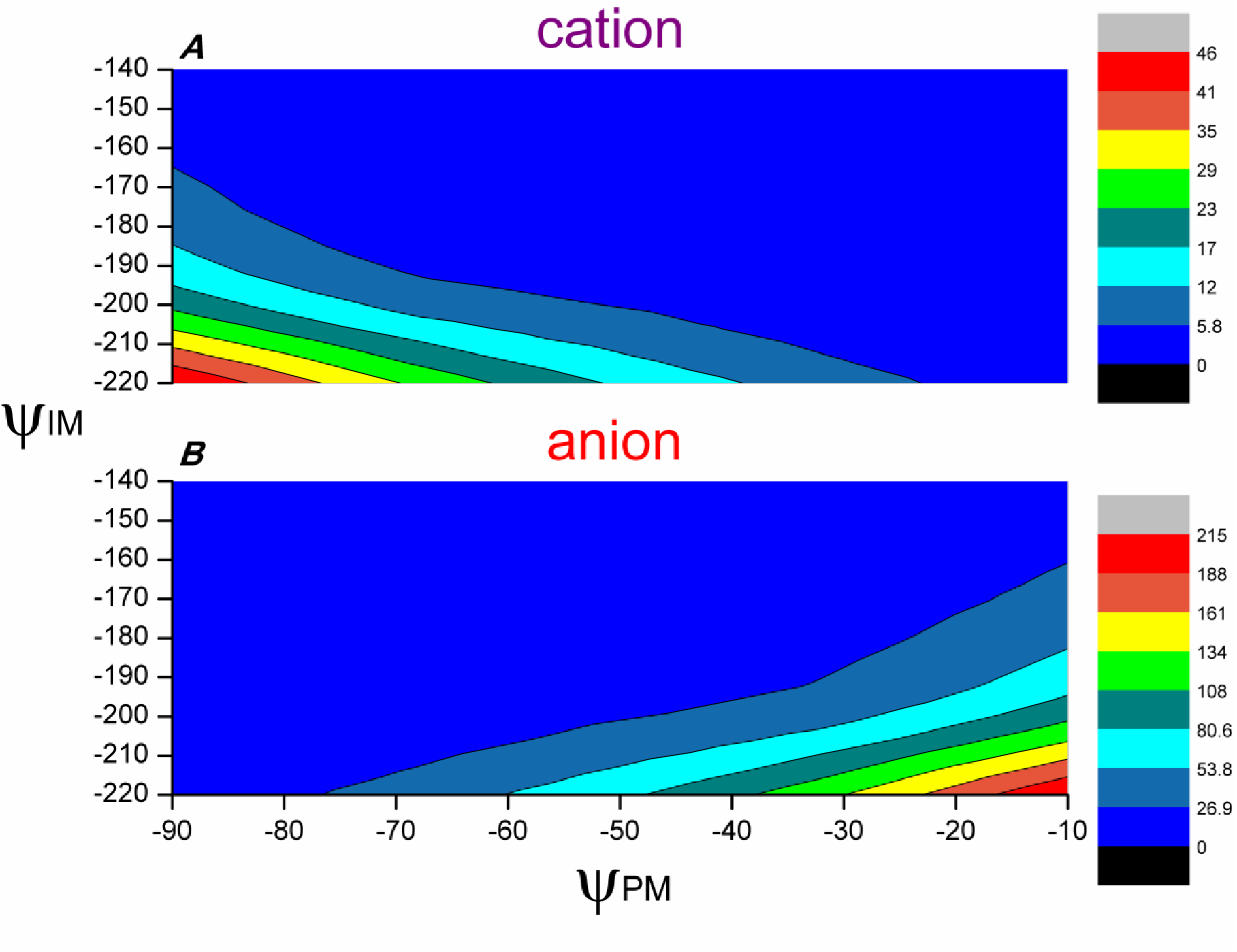
Exploring model parameter space. (*A*) The *Ratio* of cation accumulation in the mitochondrial matrix of a cancer cell (vs. normal cell) as cancer Ψ_PM_ and Ψ_IM_ varies (Ψ_OM_ = 0 mV). (*B*) The *Ratio* of anion accumulation in the IMS/cytoplasm of a cancer cell (vs. normal cell) as cancer Ψ_PM_ and Ψ_IM_ varies (Ψ_OM_ = 0 mV). Note that the scale keys of the two panels are different.

Peak cation accumulation in the matrix, over and above that of a normal cell, is when cancer Ψ_PM_ and Ψ_IM_ are both hyperpolarised. Peak anion accumulation in the IMS (or cytoplasm), over and above that of a normal cell, is when Ψ_PM_ is depolarised and Ψ_IM_ is hyperpolarised. The latter aligns with the reality of cancer cells, which have a depolarised Ψ_PM_ [4-10] and a hyperpolarised Ψ_IM_ [11-23]. So, the model indicates that lipophilic anions, rather than cations, are better as anti-cancer medicines. Note that the scale keys of the two panels are different and that the anion case has much larger numbers.

### Model V1 and Ψ_OM_

Figures 1, 2 and 3 considered normal and cancer cells to have equal Ψ_OM_ (=0). Figure 4 shows that the lipophilic cation/anion uptake differential between normal and cancer cells (Model V1) is modified if cancer cells have a different Ψ_OM_ value than normal cells. It shows how the *Ratio* in accumulation, between normal and cancer cells, changes with Ψ_DIFF_ = (cancer Ψ_OM_) − (normal Ψ_OM_). Panels *A* and *B* relate to cations/anions, panels *C* and *D* relate to di-cations/di-anions. Panels on the left present whole cell accumulation (cytoplasm + IMS + matrix), panels on the right present accumulation in a specific cellular compartment: in the mitochondrial matrix for cations/di-cations and in the mitochondrial IMS for anions/dianions.

**Figure 4.**
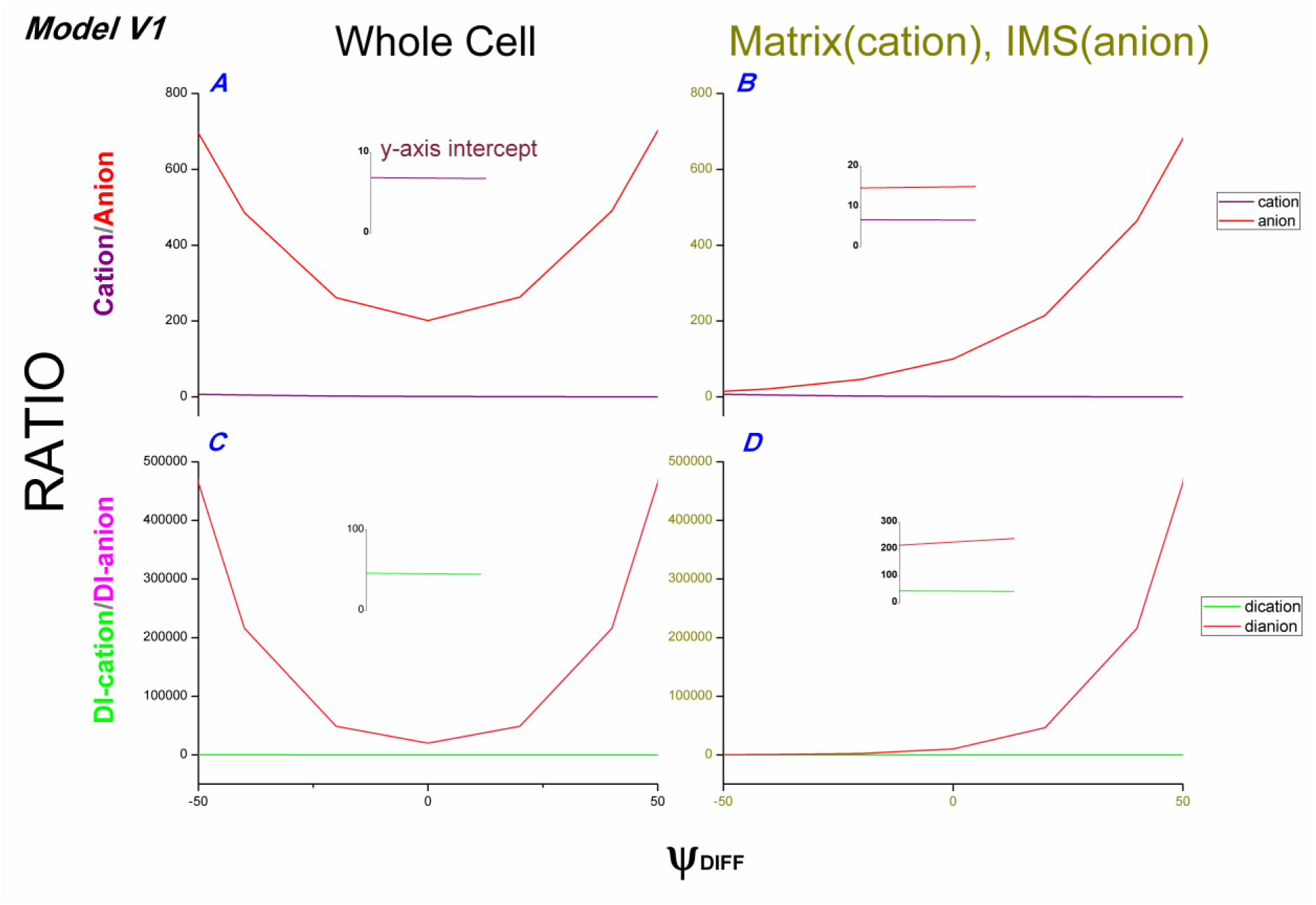
Model V1 output as Ψ_OM_ is varied. x-axis for all panels is Ψ_DIFF_ (= cancer Ψ_OM_ – normal Ψ_OM_). (*A*) Cation/anion accumulation *Ratio* (cancer vs. normal) in the whole cell. Inset graph shows where the cation plot intercepts the y-axis. (*B*) Cation accumulation *Ratio* (cancer vs. normal) in mitochondrial matrix (purple plot); anion accumulation *Ratio* in mitochondrial intermembrane space (IMS, red plot). Inset graph shows where both plots intercept the y-axis. (*C*) Di-cation/di-anion accumulation *Ratio* (cancer vs. normal) in the whole cell. Inset graph shows where the di-cation plot intercepts the y-axis. (*D*) Di-cation accumulation *Ratio* (cancer vs. normal) in mitochondrial matrix; di-anion accumulation *Ratio* in IMS. Inset graph shows where both plots intercept the y-axis.

A negative Ψ_DIFF_ increases the DLC uptake differential (*Ratio*) between cancer and normal cells at their mitochondrial matrix (*B*, *D*), which increases the *Ratio* for whole cell DLC accumulation (e.g. Ψ_DIFF_ = −50: *Ratio* = 7 for cations, 46 for dications; *A*, *C*). But a positive Ψ_DIFF_ decreases the DLC uptake differential between cancer and normal cells at their mitochondrial matrix (*B*, *D*) and for the whole cell (e.g. Ψ_DIFF_ = +50: *Ratio* = 0.2 for cations, 0.02 for dications; *A*, *C*). So, in this latter case, normal cells actually uptake more DLC than cancer cells.

Both a positive or negative Ψ_DIFF_ increases the lipophilic anion/di-anion accumulation by cancer cells as compared to normal cells, when considering whole cell accumulation. A negative Ψ_DIFF_ decreases the accumulation in the IMS (e.g. Ψ_DIFF_ = −50, *Ratio* in IMS = 15 for anions, 215 for di-anions; *B*, *D*) but it increases the accumulation in the cytoplasm, to confer a massive *net* overall increase in whole cell accumulation (e.g. Ψ_DIFF_ = −50, *Ratio* in whole cell = 696 for anions, 464,375 for di-anions; *A*, *C*). A positive Ψ_DIFF_ decreases the accumulation in the cytoplasm, but increases the accumulation in the IMS (e.g. Ψ_DIFF_ = +50, *Ratio* in IMS = 681 for anions, 464,159 for di-anions; *B*, *D*), to confer a massive *net* overall increase in whole cell accumulation (e.g. Ψ_DIFF_ = +50, *Ratio* in whole cell = 703 for anions, 464,421 for di-anions; *A*, *C*).

Please take note that the anionic approach is significantly strengthened, if Ψ_DIFF_ is either positive or negative, and is extant if Ψ_DIFF_ = 0 (*A*, *C*). Whether the real Ψ_DIFF_ is positive or negative will dictate whether we use lipophilic anion drugs to target molecules in the cytoplasm or IMS. If Ψ_DIFF_ = 0, then targets in the IMS and cytoplasm are equally valid. At present, there is no evidence – but there is a theory that – Ψ_OM_ varies between normal and cancer cells and that Ψ_DIFF_ is positive (refer *Appendix*). In this case, targets in the IMS are better. Anionic protonophores (uncouplers) are an exceptionally good drug class to target to the IMS of cancer cells.

### Model V2 and Ψ_OM_

Figure 5 follows the same format as Figure 4, but uses Model V2 as the cancer cell instead of Model V1. The same trends are seen, although the numbers are different.

**Figure 5.**
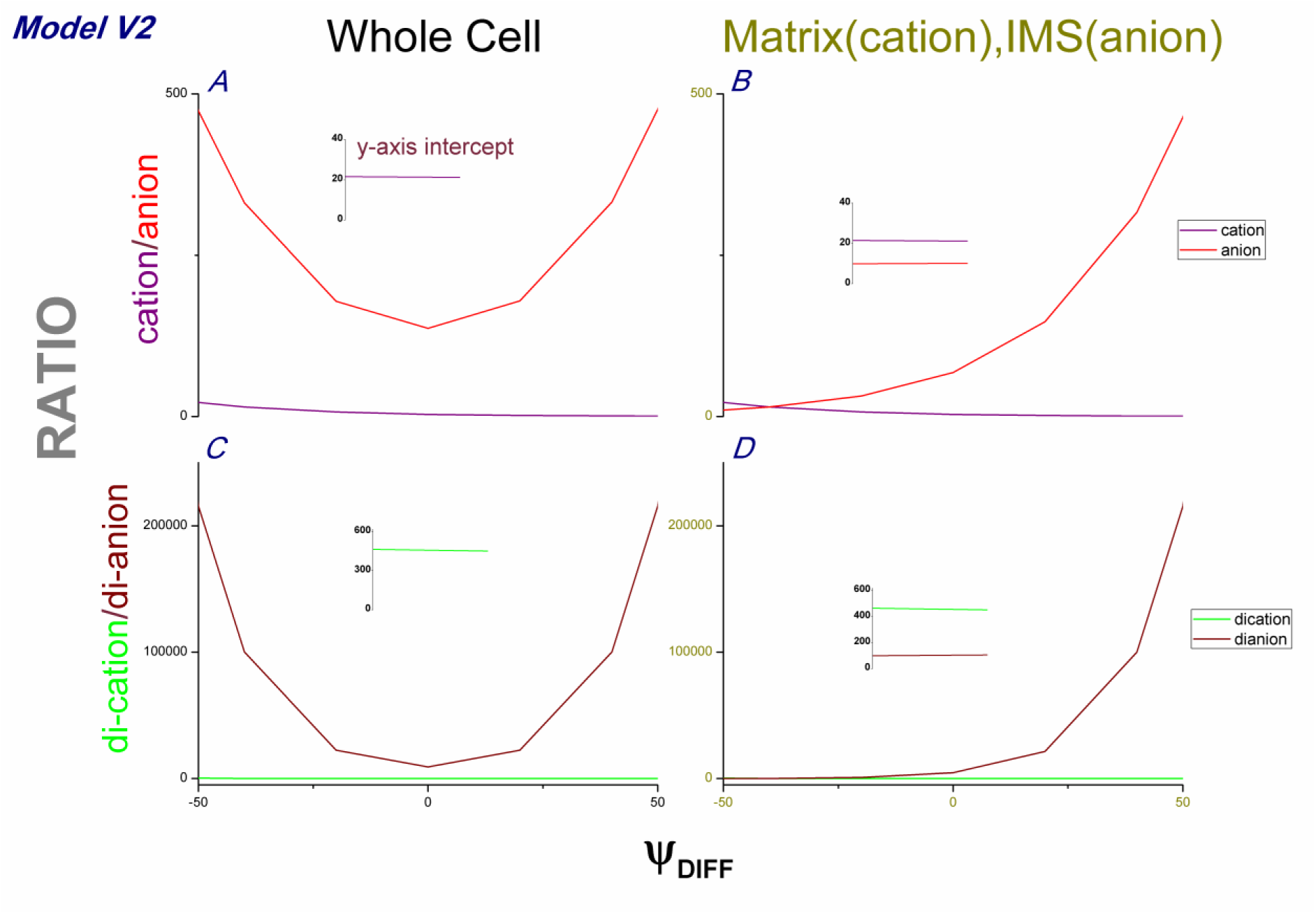
Model V2 output as Ψ_OM_ is varied. x-axis for all panels is Ψ_DIFF_ (= cancer Ψ_OM_ – normal Ψ_OM_). (*A*) Cation/anion accumulation *Ratio* (cancer vs. normal) in the whole cell. Inset graph shows where the cation plot intercepts the y-axis. (*B*) Cation accumulation *Ratio* (cancer vs. normal) in mitochondrial matrix (purple plot); anion accumulation *Ratio* in mitochondrial intermembrane space (IMS, red plot). Inset graph shows where both plots intercept the y-axis. (*C*) Di-cation/di-anion accumulation *Ratio* (cancer vs. normal) in the whole cell. Inset graph shows where the di-cation plot intercepts the y-axis. (*D*) Di-cation accumulation *Ratio* (cancer vs. normal) in mitochondrial matrix; di-anion accumulation *Ratio* in IMS. Inset graph shows where both plots intercept the y-axis.

A negative Ψ_DIFF_ increases the DLC uptake differential (*Ratio*) between cancer and normal cells at their mitochondrial matrix (*B*, *D*), which increases the *Ratio* for whole cell DLC accumulation (e.g. Ψ_DIFF_ = −50: *Ratio* = 22 for cations, 464 for dications; *A*, *C*). But a positive Ψ_DIFF_ decreases the DLC uptake differential between cancer and normal cells at their mitochondrial matrix (*B*, *D*) and for the whole cell (e.g. Ψ_DIFF_ = +50: *Ratio* = 0.6 for cations, 0.2 for dications; *A*, *C*). So, in this latter case, normal cells actually uptake more DLC than cancer cells.

Both a positive or negative Ψ_DIFF_ increases the lipophilic anion/di-anion accumulation by cancer cells as compared to normal cells, when considering whole cell accumulation. A negative Ψ_DIFF_ decreases the accumulation in the IMS (e.g. Ψ_DIFF_ = −50, *Ratio* in IMS = 10 for anions, 100 for di-anions; *B*, *D*) but it increases the accumulation in the cytoplasm, to confer a massive *net* overall increase in whole cell accumulation (e.g. Ψ_DIFF_ = −50, *Ratio* in whole cell = 474 for anions, 215,544 for di-anions; *A*, *C*). A positive Ψ_DIFF_ decreases the accumulation in the cytoplasm, but increases the accumulation in the IMS (e.g. Ψ_DIFF_ = +50, *Ratio* in IMS = 464 for anions, 215,444 for di-anions; *B*, *D*), to confer a massive *net* overall increase in whole cell accumulation (e.g. Ψ_DIFF_ = +50, *Ratio* in whole cell = 476 for anions, 215,548 for di-anions; *A*, *C*).

### Whether Ψ_DIFF_ is positive or negative (rather than 0) will dictate whether lipophilic anions are deployed against targets in the cytoplasm or IMS

Figure 6 shows how anion/cation accumulation (of charge number 1-10) varies between cancer/normal cells (via a Ratio comparison value) when Ψ_DIFF_ is changed, according to Model V1 (A) and V2 (B). Both models show largely the same trends, but their magnitude is greater with V1 in panel *A*. Anionic accumulation Ratio (red lines) is consistently greater than cationic (green lines), no matter the charge number or Ψ_DIFF_ value. When Ψ_DIFF_ = +50, the cationic accumulation in normal cells is actually greater than in cancer cells (Ratio = negative). So, no therapeutic margin in this case! When Ψ_DIFF_ = 0, it is equal for Model V1: so no therapeutic margin here either. However, cationic does fare better when Ψ_DIFF_ = −50, for both V1 and V2, or when Ψ_DIFF_ = 0 (with Model V2 only). The Ratio is extremely substantial for anionic at all data points. Note how there is equal anion accumulation Ratio when Ψ_DIFF_ = −50 or +50. However, in the former case, the accumulation for the cytoplasm is given and in the latter case, the IMS. So, whether Ψ_DIFF_ is −50 or +50, rather than 0, does not compromise the anionic therapeutic case. In fact, it enhances it. But it will dictate whether it is used to target molecules or processes in the cytoplasm or IMS. If Ψ_DIFF_ = 0, targets in the cytoplasm or IMS are equally valid, and still extremely amenable, to lipophilic anion therapy. Note the substantial values upon the y-axis; particularly for anionic cases.

**Figure 6.**
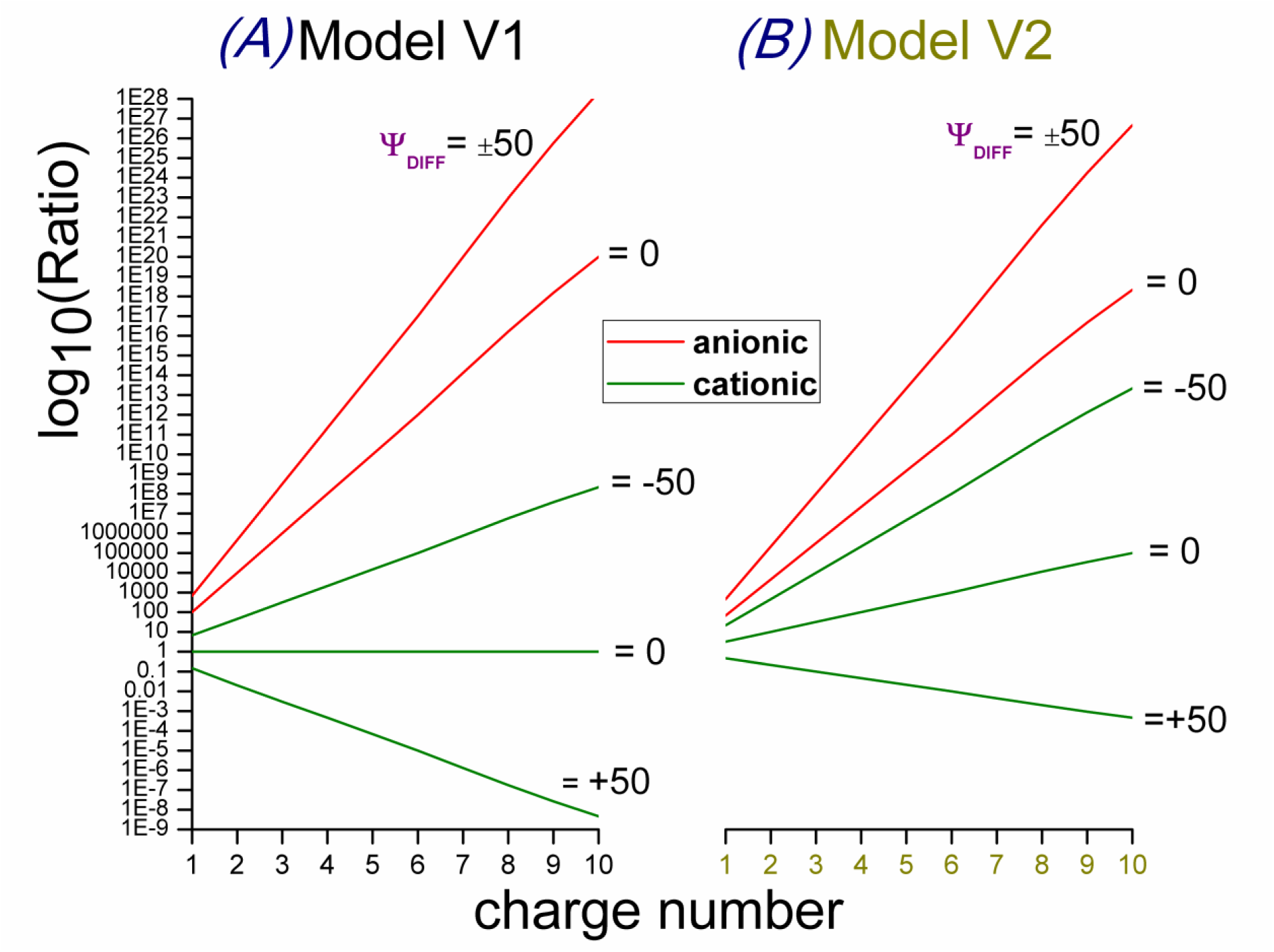
The differential in anion/cation accumulation, between cancer and normal cells (*Ratio*), changes with charge number *and* Ψ_DIFF_; where Ψ_DIFF_ (= cancer Ψ_OM_ – normal Ψ_OM_). x-axis presents charge number, y-axis presents log10 of *Ratio:* the differential in accumulation between cancer and normal cells. Different plots represent different Ψ_DIFF_ values (as labelled) and/or a cationic/anionic distinction. Anionic plots are red, cationic plots are blue. (*A*) Model V1 output. (*B*) Model V2 output.

### Model assumption

The model assumes that the cell’s membrane potentials are fixed. This may well be the case. However, as the anionic/cationic molecules assort, according to the membrane potentials, they will exert forces that act against the maintenance of these potentials, which will make the cell have to work harder to maintain them. For example, the accumulation of anions in the mitochondrial intermembrane space, which is largely driven by Ψ_IM_, is in turn a depolarising/eroding force to Ψ_IM_. So, the cell will have to take compensatory action to maintain Ψ_IM_ e.g. pumping more protons across the IM. The force exerted across the IM by these lipophilic anions is given by the Nernst equation, with its intermembrane and matrix concentrations as inputs. It returns a voltage that this anionic assortment, across the IM, is exerting upon the membrane potential. Lipophilic anions exert a hyperpolarising force across the PM and a depolarising force across the IM; more so for cancer than normal cells. Both these forces are detrimental to cancer cells. Depolarisation across the IM undermines the proton motive force and moves the cell towards apoptosis – if cells can’t maintain a hyperpolarised Ψ_IM_, the voltage-dependent permeability transition pore opens and drives programmed cell death [65]. Hyperpolarisation across the PM stops cancer cells from proliferating: a depolarised Ψ_PM_ is necessary for cancer cells (causative to cancer, not merely correlative [4-5, 7]). So, if lipophilic anions can move these membrane potentials, this could bring therapy. In a manner completely non-reliant upon any “lock and key” molecular interaction, which cancer cells can mutate their way to resistance from. The forces exerted don’t change with the charge number of the anion: because a larger charge number doesn’t change the ratios of concentration across the membranes, which is the salient numeration, not the absolute concentrations. Nor do they change with user-inputted, extracellular concentration in the model for the same reason. The added potential driven by the lipophilic anions at the PM and IM, for normal cells, is −70 mV and +280 mV respectively. For cancer cells (Model V1), it is −190 mV and +400 mV respectively. So, once again, we see lipophilic anions acting disproportionally against cancer cells.

### Informatics

In the *Supplementary Information* there is a list of anionic compounds, from the Pubchem database [66] (as of Dec 2015), that conform to Lipinski’s rule of 5 [58] (as specified by Pubchem) *and* that have had some prior bioactivity established. I propose that the most anionic – especially those that co-correspond with the most promising bioactivity – should be tested first. Very anionic compounds that inhibit/disrupt some key mammalian, cellular process should be prioritised. If the molecule is anionic enough, normal cells should be safe within the therapeutic margin. Anionic protonophores (uncoupling drugs) are a particularly exciting prospect and I deal with these specifically in a further, forthcoming paper. In the *Supplementary Information*, I also list some further molecules, with established bioactivity, that conform to some Lipinski’s rule variants e.g. where the mass cut-off is 600, rather than 500 Daltons, and the hydrophobicity measure (*logP*) is slanted towards more hydrophobic (1 to 7). Of course, these lists of molecules are not exhaustive. For example, there are many anionic molecules in the Pubchem database that are yet to be tested for bioactivity, or a relevant bioactivity, and many more molecules that are yet to be found and included in this database. But it is useful as an initial prioritising guide for anti-cancer tests. My aim is that molecules can be tested systematically, within a strong conceptual framework, rather than randomly. I propose that anionic character permits an anti-cancer drug to target machinery that both normal and cancer cells rely on and increases the efficacy of drugs targeting cancer specific processes. My co-use of Lipinski’s rule of 5 can be modified, relaxed or dropped.

## DISCUSSION

### The model valuably reconciles a divergent DLC literature

There is a mysterious dichotomy in the literature. DLCs have shown such promise in pre-clinical studies (*in vitro* and mouse) [11-16, 20, 24-40] and yet no DLC has passed clinical trials to date [41-43]. How can this be explained? Well, it has actually been known since the 1980s that DLCs won’t work for all cancers [31]. Furthermore, that the successes come with cancer cells accumulating more of a DLC than normal cells and the failures relate to cancer not accumulating/retaining the DLC more than normal cells [31]. Indeed, the DLCs employed thus far don’t tend to target processes unique to cancers but are poisonous to normal cells also [15] e.g. rhodamine 123 inhibits ATP synthase [36]. The DLC approach, to date, simply relies upon a higher DLC accumulation in cancers. What is the basis to this selective accumulation, and thence action?

My model valuably reconciles why DLCs work in some cases and not others. It shows why some cancer types will accumulate a DLC more and be more susceptible than normal cells (Model V2), and others won’t (Model V1). I propose that the positive DLC experimental studies are upon Model V2 cancers, the negative clinical DLC studies upon Model V1 cancers. The model is an insightful abstraction, with its foundation in the fidelity of the Nernst equation and parameterisation to experimental data. With this new insight, I hope that the DLC approach to cancer therapy can be revisited.

### Multidrug resistance

There is a pertinent factor not considered in the model, but discussed here. Different cancer cell lines can differ in their expression of DLC efflux mechanisms e.g. the *p*-glycoprotein efflux pump (*p*-gp, MDR1 gene product). This is a multidrug resistance protein that pumps out positive lipophilic compounds e.g. rhodamine 123 [67]. Some have suggested that differential *p*-gp expression is an additional reason why DLCs can kill some cancer cell lines, at concentrations that don’t kill normal cells, and not others [68-69]. Incorporating this into my model: if a cancer is Model V1 type it will be comparatively (vs. normal cell) DLC resistant, whether it expresses *p*-gp or not. If a cancer is Model V2 type, but expresses *p*-gp, it will be comparatively DLC resistant. If a cancer is Model V2 type, but doesn’t express *p*-gp (or if this is pharmacologically inhibited or genetically knocked down), it will be comparatively DLC susceptible. DLCs can be pumped out of cells by other multidrug efflux proteins also e.g. rhodamine 123 is a substrate for MRP1 [67]. But this can be inhibited in addition, as required. Numerous drugs inhibit *p*-gp and/or MRP1 [70-72]). So, DLC therapy may be assisted, in some cases, by co-therapy with these inhibitors.

### Interpreting specific DLC studies in the light of the biophysical model

Some DLC studies have compared human cancer cell lines with a normal African green monkey kidney epithelial cell line, CV-1 [16]. Or with normal kidney epithelial cells from other non-human species [13, 20, 30]. So, the cancer cell lines tested don’t correspond in origin to any of the normal tissues tested. This adds an unwanted variable: the cell type (and species). It matters because different cell types can have different *p*-gp expression levels. For example, CV-1 expresses *p*-gp whereas many of the cancer cell lines tested in these studies do not e.g. MCF-7 doesn’t [73]. Active DLC efflux by *p*-gp in CV-1 cells, and no such efflux in the cancer cell lines lacking *p*-gp, might cause the observed differential in DLC accumulation and action. Wherein the DLC kills cancer cells and not normal CV-1 cells. Indeed, if *p*-gp is inhibited by verapamil in CV-1 cells, they accumulate and become susceptible to DLCs also [73]. In contention to this account, [40] report that their CV-1 cells do not express *p*-gp (because of a susceptibility to adriamycin).

There are cases in the DLC literature where a cancer cell is compared to directly correspondent normal cells. Mouse or rabbit bladder epithelial cells were transformed using a carcinogen, and the transformed cells were compared with the original, untransformed cells [12]. In this case, differential *p*-gp expression is less likely as the cancer cells probably inherit and express the same *p*-gp level as the normal cells. In this controlled case (for cell type and *p*-gp expression) the cancer cells do accumulate more rhodamine 123 than normal cells. So, these cancer cells align with Model V2 rather than V1. However, one of these derivative cancer cell lines doesn’t accumulate more DLC and I would align it instead with Model V1 (or alternatively it has acquired a greater *p*-gp expression). In another controlled study, no preferential accumulation or toxicity of rhodamine 123 was found in breast cancer cells compared to normal breast cells [73]. I would equate these cancer cells to Model V1.

[11] compares rhodamine 123 accumulation and retention in a mouse bladder epithelial cancer cell line (MB 49) to normal mouse bladder epithelial cells. Ensuing differences in colony formation are studied. It finds that rhodamine 123 impairs cancer cell colony formation as compared to normal cells, probably as a function of their greater DLC accumulation and retention. I suggest these cancer cells align with Model V2.

Rhodamine 123 accumulation in malignant and non-malignant breast epithelial cell lines was compared. [38]. 100% of the non-malignant cell lines showed weak accumulation. ~71% of malignant cell lines showed strong accumulation, ~21% of malignant cell lines showed weak accumulation and ~7% of malignant cell lines had cellular heterogeneity: some cells accumulated strongly, others weakly. The strong accumulators equate to Model V2, the weak accumulators to some model intermediate between Models V1 and V2 (note that Models V1 and V2 are discrete, binary abstractions that actually represent a continuum of diversity within bounds).

The MIP101 cancer cell line is resistant to rhodamine 123, a DLC, and yet has a hyperpolarised Ψ_IM_ like the CX-1 cancer cell line, which is susceptible to it [36]. A relevant distinction is that MIP101 expresses *p*-gp and CX-1 doesn’t [74]. So, some cancers express *p*-gp and are resistant to DLC accumulation, retention and poisoning [36]. DLC resistant cancers have been reported since the first papers on DLCs in the 1980s [12-13]. [75] found ~30% of human tumours to express *p*-gp (sample size: 182). Clinical case cancers could inherit *p*-gp expression from their host tissues, as *p*-gp expression is found in many normal cells [75-76]. In cases where they don’t, they may still acquire it. A single cancer cell may acquire its expression by mutation, and then be selected for by drug exposure [77].

### The disappointment of DLCs in clinical trials to date

MKT-077 is a DLC that selectively killed cancer cells *in vitro* and in xenograft mouse models [27-28, 34, 40]. However, it failed clinical trials [41-42]. I suggest that in trialling, MKT-077 found itself confronted by Model V1 cancers (that may or may not have expressed *p*-gp) or Model V2 cancers with *p*-gp expression. And was found wanting. In this case, the cancer cells don’t accumulate any more of the DLC than key normal cells and so there is no therapeutic margin. Indeed, normal cells are affected at doses required to kill cancer cells e.g. kidney cells. MKT-077 failed two independent phase 1 trials because it caused renal toxicity [41-42]. MKT-077 also caused renal toxicity in toxicological evaluations in animals [42] and accumulated most in the kidney cortex [78]. But it caused liver, rather than kidney, problems in a different animal toxicology evaluation [28]. The DLCs, dequalinium and rhodamine 123, also showed a lot of promise *in vitro* and *in vivo* (nude mice & rats) [24, 30, 31-32].

However, in phase 1 trials rhodamine 123 couldn’t kill cancer cells, to the stringency of statistical significance, at the maximally tolerated dose [43]. Although it did not cause nephrotoxicity and renal magnesium wasting like MKT-077. Its point of failure was different. This is despite rhodamine 123 being accumulated extensively by rat kidney cells, *ex vivo*, in an isolated kidney [79]. Dequalinium damaged the kidneys (and liver) of mice in toxicological evaluations that explored maximal dosages permissible [80]. In an *Appendix* to this paper I discuss in some detail why I think MKT-077 causes kidney damage.

### The future of DLC therapy

F16 [14], AA1 [25] and gallic acid derivatives [29] are other DLCs that have showed *in vitro* and *in vivo* promise with cancer cell lines. However, they haven’t been pursued in clinical trials to date, to my knowledge. I suggest that this is because the DLC approach to cancer, that once showed such promise, is now in crisis. The failure of MKT-077 and rhodamine 123 in clinical trials has bought doubt. I don’t know why. It has been known since the 1980s that DLCs won’t work for all cancers [31]. And that these failing cases are related to the cancer cell lines not selectively accumulating/retaining the DLC more than normal cells. I think it was maybe thought that greater DLC accumulation/retention was a widespread feature of cancerous epithelial cells, as these were shown most amenable to DLC treatment in preclinical trials e.g. [31]. So, if a patient had a cancer epithelial in origin, then they were chosen as fit to incorporate into the trial (in [41] all patients had cancer of epithelial origin, for [42] it’s unclear). This was an assumption though, which I suggest is misplaced. I propose that DLC susceptibility doesn’t scale in such a neat, tidy assortment. And that in any future DLC clinical trials, patients should be selected on the basis of tumour biopsies. Investigated *in vitro* to see if their cancer can selectively accumulate/retain the DLC. With the emergent trend for “personalised medicine”, this may be more conceivable than it has been in the past.

### Lipophilic anions rather than cations

The rationale for DLC therapy has been that cancer cells have a more hyperpolarised Ψ_PM_ *and* Ψ_IM_ [16, 20]. But I argue that cancer cells actually have a more depolarised Ψ_PM_ [4-10] and a more hyperpolarised Ψ_IM_ [11-23]. I show that leveraging differentials in Ψ between cancer and normal cells, with charged lipophilic molecules, is still a valid therapeutic tact. And that this can be with cations, in certain cancer cases, as aforementioned. ***But*** that ideally this should be with anions rather than cations. Targeting molecular species and processes in the cytoplasm, and/or mitochondrial intermembrane space, rather than the mitochondrial matrix. I propose the use of delocalised lipophilic anions (DLAs); DLAs rather than DLCs. My model predicts that DLAs can have a universal utility, against all cancers, where DLCs have a delimited utility.

For cancer cells: their stereotypical depolarisation in Ψ_PM_, and hyperpolarisation in Ψ_IM_, decreases and increases DLC accumulation respectively. These cancel and render little or no DLC accumulation differential from normal cells. However, for DLA accumulation, these Ψ_PM_ and Ψ_IM_ changes are additive rather than subtractive. This compounding causes a substantial differential in DLA concentration from normal cells; a significant therapeutic margin.

The immense magnitude of this differential accumulation suggests that even molecular species that both cancer and normal cells rely on can be targeted. On the basis that much more of the drug will accumulate in cancer than normal cells. So, cancer cells will be killed at doses that don’t kill normal cells. There is a rich zoo of cytoplasmic/IMS molecules; thousands of different molecular species. The model predicts that they are all good targets for anti-cancer medicines, *if* the drug is a lipophilic anion. This is the opening of a new front in the war against cancer.

### Multidrug pumps

Active efflux could be a conceivable route to resistance against lipophilic anion drugs. Indeed, a problem with DLC therapy can be that DLCs are recognised and pumped out by ABC multidrug transporters e.g. *p*-gp [67]. While *p*-gp favours efflux of lipophilic cations, the nine members of the MRP family favour efflux of lipophilic anions [81]. Although this charge distinction isn’t complete e.g. MRP1 effluxes DLCs also e.g. rhodamine 123 [67]. Breast cancer resistance protein (BCRP) effluxes both positive and negative molecules [82]. However, a distinction must be made between the use of lipophilic anions rather than cations. My model shows that the driving force for anions into cancer cells, as compared to normal cells, is massive. It can be on the scale of hundreds, thousands, millions, billions, trillions etc. By contrast, for cations it is slight to non-existent. So, cancer cells would have to acquire a massive anion efflux capability, over and above that of normal cells, to outweigh the massive driving forces for lipophilic anion entry, which they are subjected to well in excess of normal cells. Furthermore, lipophilic anion extrusion comes at a cost. It requires ATP hydrolysis and large lipophilic anion influxes, and subsequent effluxes, could cause a cellular ATP crisis in cancer cells. This could bring therapy in and of itself. It could actually kill cancer cells that use multidrug resistance (MRP) pumps and select for cancer cells without such pumps; the inverse of typical chemotherapies. So, pre-treatment with lipophilic anions could kill cancer cells with multidrug resistance pumps (of MRP class) and leave a remainder without such pumps, but these are then susceptible to a chemotherapeutic that if used alone, without this pre-treatment, would merely be pumped out of the cell and rendered impotent. So, we are confronting the problem of multidrug resistant cancer, which is typically a death sentence. Alternatively to pre-treatment, lipophilic anion therapy could be used in co-treatment with chemotherapy to flood the efflux system, undermining its extrusion of the chemotherapeutic, and bring an ATP crisis that potentiates the kill action of the chemotherapy. Lipophilic anion entry requires ATP to counteract, and extrude the anion, but at the same time this entry undermines ATP generation. The accumulation of the lipophilic anion in the IMS is a depolarising force to Ψ_IM_, which acts to reduce the proton motive force. So, concurrent with expending ATP (for their extrusion), lipophilic anions act to undermine ATP generation also. This is true of any lipophilic anion; but even more so if the molecule is a protonophore/uncoupler. In this case, their compromise of ATP generation may render them efflux resistant, given that their extrusion hinges upon ATP. This is all done without requiring any “lock and key” interaction, which is easy for cancer cells to mutate out of and attain resistance. Furthermore, anions can be pumped out in a complex with glutathione (GSH) [83]. So, in this case, for every anion molecule pumped out, a glutathione molecule is lost. Glutathione is integral to the ROS mitigation system. This mitigation system is absolutely fundamental to cancer cells: much of their danger comes from their immortality, which in turn comes from their lesser ROS generation and greater ROS mitigation (elaborated on later). So, lipophilic anions will attack a pillar of this immortality and thence undermine cancer cell pathology. Many chemotherapies and radiotherapies kill cancer cells by ROS generation and lipophilic anion co-treatment will potentiate their effects. This is valuable as these methods are typically unspecific for cancer, and generate ROS across normal and cancer cells. But when combined with lipophilic anion therapy, which is very selective for cancer cells, greater specificity can be attained. Lipophilic anion therapy will reduce the ROS levels required to kill cancer cells, which will thus reduce the collateral ROS generation/damage in normal cells. Glutathione is typically more abundant in cancer cells, such is its importance to them [84], and lipophilic anions may sabotage at this point of reliance.

### A depolarised Ψ_PM_ and hyperpolarised Ψ_IM_ are integral to the metabolic program employed by cancer cells

Aerobic respiration is O_2_ dependent and uses glycolysis, the Krebs cycle and oxidative phosphorylation (OXPHOS) to produce ATP [85-87]. Aerobic glycolysis is the sole use of glycolysis to produce ATP, even in the presence of O_2_ [21-22, 88-107].

Cancer cells have a depolarised Ψ_PM_ (~-10 mV) and a hyperpolarised Ψ_IM_ (~-200 mV) compared to normal cells (Ψ_PM_ = ~-70 mV, Ψ_IM_ = ~-140 mV) [4-10, 11-23]. A route to resistance from lipophilic anions would be for the cancer cell to re-align its Ψ_PM_ and Ψ_IM_ values to that of a normal cell. So reduce its drug accumulation to the level of a normal cell, and remove the therapeutic margin. However, I propose that these altered Ψ_PM_ and Ψ_IM_ values are integral to, and a hallmark of, the metabolic program of cancer. Indeed, I propose that cancer cell metabolism is similar to that of embryonic stem (ES) cells. And that the pathology of cancer is that cells, as a result of mutations, switch into this physiological metabolic program, but at an inappropriate time which confers pathology. Indeed, they share genetic expression fingerprints [108-109] and ES cells have a hyperpolarised Ψ_IM_ [110] and depolarised Ψ_PM_ also [7-10]. They both employ aerobic glycolysis some or all of the time [21-22, 88-107, 111], are immortal (divide forever without limit) [112-113] (as a function of using aerobic glycolysis [114]), respond to ROS damage by apoptosis rather than repair [22, 115] and can proliferate rapidly.

### Normal cells are mortal

Normal adult cells cannot divide forever [85-87]. They utilise OXPHOS, which increases their ATP yield from glucose, but that also produces reactive oxygen species (ROS). These ROS cause DNA damage (ageing) [116]. Eventually, this damage accumulates to the point that the information fidelity of the cell is damaged so much that it cannot replicate itself further. Normal adult cells are programmed to limit their number of replications (“telomere clock”, Hayflick limit, 50-70 divisions [85-86]) so that they have a certain number of divisions by design, rather than reaching a later limit because of damage. So, a normal adult cell cannot divide forever, without limit. But a cancer cell can because – I would argue – it utilises a different metabolism. Cancer cells shunt OXPHOS, and use aerobic glycolysis, some or all of the time [21-22, 88-107]. As I’ve suggested previously [102], at the very least during S phase: when DNA is unwound from protective chromatin, exposed and vulnerable as it is being replicated. Especially because when negative DNA and positive histones separate their charges are less screened and neutralised. The non-sequestered positive charges can assort to attract more superoxide (O_2_•−) where it is more likely to interact with, and damage, negative DNA. Especially, when the positive histones – perhaps with O_2_•− in electrostatically attracted proximity – return to re-complex with DNA.

### Aerobic glycolysis, a hallmark of cancer, conveys immortality, a further hallmark of cancer

I argue that cancer cells reduce ROS at source and sink by using aerobic glycolysis, some or all of the time, which conveys them immortality. Firstly, OXPHOS is shunted, by shunting NADH production [102], which reduces ROS generation. Secondly, ROS mitigation is upregulated by increasing NADPH production. So, as compared to normal cells, cancer cells decrease [NADH] and increase [NADPH]. Higher NADPH in cancer cells has been observed by spectroscopy [117]. Higher NADH has been reported also [118] but I think the authors misinterpret their data and are actually observing higher NADPH (the spectroscopy used in this latter study can’t distinguish between NADH and NADPH). The elevated glycolysis of cancer cells protects from oxidative damage [114]. High glycolytic rate permits a high flux into the pentose phosphate pathway (PPP), which branches from glycolysis. It produces NADPH from NADP+, which is needed for glutathione (GSH)-dependent anti-oxidant mechanisms. To protect, GSH needs to be in its reduced form and NADPH puts it into this reduced form (as it is converted to NADP+). GSH is needed by glutathione peroxidase (GP), which converts hydrogen peroxide (a ROS) into water. Upstream, superoxide dismutase (SOD) converts superoxide (O_2_•−, a ROS) into hydrogen peroxide. Increased GP activity will pull through greater SOD activity.

With these lower ROS levels, at key proliferation stages, cancer cells can limit their DNA damage, maintain information fidelity and divide forever without limit. So, aerobic glycolysis conveys immortality which conveys the danger of cancer. For a cancer to be truly dangerous it needs to be immortal (divide without limit) or at least by able to divide enough times to kill the host (a lot! >> Hayflick limit). I propose that benign cancers are those that don’t tap into aerobic glycolysis sufficiently, aren’t retracted along the proliferation/differentiation continuum sufficiently, and eventually accrue sufficient damage that they can’t divide any longer. The really dangerous cancers are those that retract further back along the proliferation/differentiation continuum, to the metabolic program of ES cells, and employ aerobic glycolysis enough, as a proportion of their energy mix, to attain immortality. Indeed, there is a correlation between markers of aerobic glycolysis and a poor clinical prognosis; as I shall now discuss.

I suggest that a depolarised Ψ_PM_ and hyperpolarised Ψ_IM_ are markers of using aerobic glycolysis, and so I would argue a marker for the aggressiveness and danger of the cancer. Experimentally, when cancer cells are switched out of aerobic glycolysis, into aerobic respiration, their Ψ_IM_ is returned to that of normal cells [21-22]. The more invasive and dangerous the cancer, the more hyperpolarised its Ψ_IM_ is observed to be [17-19]. In a prior paper, I used biophysical modelling to show how and why aerobic glycolysis produces a more hyperpolarised Ψ_IM_ [23]. Here I explain why it produces a depolarised Ψ_PM_ also. Normal cells sit in neutral tissues [119-120], cancer cells reside in acidic tumours [119]. This acidity is a marker of aerobic glycolysis, which excretes H+ and lactate [121] through the monocarboxylate symporter (MCT) (a promising cancer drug target). A higher aerobic glycolysis rate produces a lower extracellular pH. The more aggressive and dangerous a cancer is, the more acidic its tumour [122]. Cancer cells must maintain their cytoplasm at neutral pH like normal cells [47-48, 119-120]. I propose that cancer cells have a depolarised Ψ_PM_ to offset their greater extracellular acidity and reduce the proton motive force (pmf) directed into the cell, in order to assist their intracellular pH homeostasis. Indeed, if a cancer cell Ψ_PM_ (~-10 mV) is hyperpolarised to a value more typical to a normal cell (e.g. ~-70 mV), its inward directed pmf is more than doubled (refer *Appendix* for calculation). So, in this way a depolarised Ψ_PM_ is an aerobic glycolysis marker. Indeed, a depolarised Ψ_PM_ is a cancer marker [4-5], with probable future applications to cancer diagnosis. Ψ_PM_ depolarisation is *required* for cancer (causative rather than merely correlative [4-5, 7) and a hyperpolarisation of Ψ_PM_ reduces tumour formation *in vivo* [5]. This finding has a therapeutic scope. Indeed, cellular entry of lipophilic anions is likely to be a hyperpolarising force to Ψ_PM_! Crucially, the Ψ_PM_ depolarisation of cancer cells alters gene expression to drive proliferation [5, 7-10]. The intermediary between Ψ_PM_ and gene expression has been experimentally established. It can be Ca^2+^ cellular entry and Ca^2+^ intracellular signalling, butyrate entry through a Na^+^-butyrate transporter (butyrate is an HDAC1 inhibitor, changing chromatin acetylation) and voltage-dependent phosphatases [7]. This list is non-exhaustive.

The predicted selective accumulation of lipophilic anions by cancer cells, because of a depolarised Ψ_PM_ and hyperpolarised Ψ_IM_, has probable applications to cancer imaging, diagnosis and therapy. Crucially, with this approach, the more dangerous the cancer, the more it is highlighted (imaging) and targeted (therapy).

### Voltage-sensitive dyes in cancer imaging

Voltage-sensitive dyes are used widely in neuroscience and cardiology for membrane potential imaging [123]. Given the aforementioned membrane potential disparities, voltage-sensitive dyes could be used, in principle, to distinguish a cancer cell from a normal cell. This might be reasonably trivial *in vitro*. In the clinical setting, this effect could be used to “light up” cancer cells during a surgery or to identify a suspicious body of cells at a body surface e.g. the skin. However, applicability is constrained by the limited tissue penetrance of the excitation/emission wavelengths for the suite of dyes that we presently have. Most work in the visual spectrum and, exceptionally, in the near infra-red [124]. Efforts should be made to develop voltage-sensitive dyes that can absorb and emit at more penetrating wavelengths. Given that cancer cells reside in acidic tumours, rather than neutral tissues [119], pH sensitive fluorescent indicators and probes may find roles in cancer diagnosis also. It is worth mentioning here that membrane potentials emit electromagnetic radiation themselves (“biophotons”) [125]. The Ψ_PM_ and Ψ_IM_ of cancer cells will emit at a different wavelength than normal cells, as a function of their different voltages, which could be leveraged in cancer diagnosis. However, these emissions are typically in the visible range. So, possibly viable for surface detection but not deep in the body interior. Although, dyes might be used as intermediaries – to absorb in this spectrum – and then to emit at a more penetrating wavelength. As Ψ_PM_ and Ψ_IM_ exert electromagnetic effects, they will in turn be modulated by electromagnetic effects, which is a potential margin for future cancer therapies.

### Neutron capture therapy

Lipinski's rule of five [58] is a trade-off to capture drugs that are hydrophilic enough to be stable and active in aqueous solution (indeed their site of action is probably in this phase e.g. in the cytoplasm or at receptors in the extracellular space) but that are also hydrophobic enough to pass through cellular membranes and so be orally bioavailable. Application of Lipinski’s rule of five tends to limit how many negative charges a molecule can have. The most negative molecule that I have found in the pubchem database [66], that still fits the rule of five, has a −6 charge e.g. [59]. So, this puts a possible upper bound on the degree of cancer targeting (at least as far as present molecules in the database). However, there are some molecules in cancer therapy that don’t need to adopt a certain, specific structure in an aqueous phase to function. They just need to be present in cancer cells more than normal cells. In this case, Lipinski’s rule of five doesn’t need to be so stringently applied and molecules can be more hydrophobic, which means they can be bigger (more carbon atoms tends to greater hydrophobicity [126]), which means they can have more negative charges. An example is boron neutron capture therapy [127]. Here the boron-10 isotope atoms do not need to be in some specific molecular configuration to bring treatment: they just need to be present; in cancer and not normal cells. So, I propose that the more negative the boron containing molecule, the better. Very large, very lipophilic, very negative molecules can be created which will yield immense levels of cancer targeting. Similar rational can be applied to gadolinium neutron capture therapy also [128]. As an example, consider a lipophilic molecule with 20 negative charges: my model predicts that cancer cells will accumulate it 10^40^ times more than normal cells (>> 1 quindecillion). As negative charges are added to a molecule it will be increasingly reluctant to enter the negative interior of a cell, but this can be offset by modifying the molecule to make it more lipophilic – perhaps by making it bigger (more carbon atoms) – which will increase its propensity to enter cells i.e. the increased anionic character can be offset by increased lipophilic character. But this offset point will be very different for normal and cancer cells, because normal cells are much more negative inside. So, molecules can be made that are incredibly admissible to cancer cells in comparison to normal cells.

### Lipophilic anions in cancer imaging and diagnosis

Lipophilic anion molecules can be used for cancer selective targeting of contrast atoms/isotopes. In order to show up tumours disproportionally in X-ray, computed tomography (CT), magnetic resonance imaging (MRI), ultrasounds, positron emission tomography (PET) and nuclear scans. For example, in mammograms that screen for breast cancer. For X-rays these lipophilic anions can be iodine, gold or barium containing compounds; for MRI: gadolinium, manganese or iron oxide containing compounds; for PET: carbon-11, nitrogen-13, oxygen-15, fluorine-18, gallium-68, zirconium-89 or rubidium-82 containing compounds. I predict that these lipophilic “contrast” anion molecules can help identify cancer earlier, decrease false-positives/false-negatives and improve clinical outcomes. Indeed, a major problem in the clinical setting can be “over-diagnosis”, which is often the treatment of a benign cancer that will never progress to malignancy [129]. For example: by unnecessary mastectomy (breast removal). This can happen because mammograms, and X-rays more generally, cannot distinguish between benign and dangerous cancers. Indeed, there is even little conceptual understanding of what is different between them. In this *Discussion*, I have offered a much-needed explanation and predict a quantitative, tractable distinction between them. That dangerous cancers have a more depolarised Ψ_PM_ and hyperpolarised Ψ_IM_, as a function of greater aerobic glycolysis use (indeed, greater blood lactate will be another marker). This means that they will accumulate lipophilic anions at much greater amounts, which can be leveraged in their diagnosis. This solution addresses one of the “Grand Challenges” identified by Cancer Research UK (with £20 million of funding available): how to distinguish benign from malignant cancer via screening mediums [129]. I envisage that my methodology will make national screening programs viable for many cancers that can’t be screened reliably with present methods e.g. lung and prostate cancer. This will catch more cancers earlier, saving lives. In this situation, the contrast molecule doesn’t have to be in any particular structure or orientation in an aqueous medium and so isn’t to be constrained by Lipinski’s rule of five. The contrast atoms/isotopes just need to be present, in any orientation to one another, in cancer cells and not normal cells. So, very large, very lipophilic, very anionic, highly cancer selective contrasting molecules can be used. This can give large signal to noise ratios.

### Tethering of lipophilic anions

A drug or molecule that isn’t lipophilic, or anionic, can be tethered to a molecule that is, which can confer upon it these properties and yield selective targeting to cancer cells (similar rational has been used previously, but with DLCs [130]). So, this strategy can be used to improve the drug complement that we already have for cancer. And to reignite some candidate drugs that may have fallen short in some regards. It can also be used to deliver photosensitizer molecules selectively to cancer cells, for photodynamic cancer therapy [131], or fluorescent molecules for cancer imaging e.g. for real time imaging of cancer during surgery. Tethering molecule candidates include the lipophilic tetraphenylborate anion (TPB) or a dipicrylamine anion (DPA) [87]. Relevantly, 8-hydroxypyrene-1,3,6-trisulfonate is an anionic (-3), lipophilic fluorophore [132].

### Nanomedicines

Thus far we have discussed how charged lipophilic molecules (cations or anions) can be selectively targeted to cancer cells. Well, the different Ψ_PM_ and Ψ_IM_ values of cancer cells may be used to disproportionally target non-lipophilic molecules to cancer cells also. When they are enclosed in a liposome (or micelle) delivery system [133] that has an associated charge, either because some or all of the molecular or ion constituents have a charge, or molecules or ions of charge are embedded in the membrane of the liposome or because a transmembrane potential is attributed to the liposome; or to smaller, constituent liposomes contained within. In parallel to the situation with lipophilic molecules, I think negatively charged liposome systems will convey much better cancer targeting than positively charged. The liposome payload can be a nutrient, metabolite, drug, molecule, antibody, DNA, plasmid, RNA, siRNA, shRNA etc. Crucially, this liposome method may deliver macromolecules at a higher concentration to cancer cells than normal cells. This relates to another of the six “Grand Challenges” identified by Cancer Research UK: to “Deliver biologically active macromolecules to any and all cells in the body” [129]. Adhering to the call, I envisage my system will deliver to all cells of the body. However, I don’t apologise if it, as I anticipate, delivers extremely disproportionally to cancer cells. The power of this postulated nanomedicine is how generic it is: any given macromolecule can be disproportionally delivered to cancer cells; a new therapeutic paradigm.

### The importance of theory

There are 10^60^ possible “drug-like” molecules (smaller than 500 Daltons) [134]. A cure for cancer is likely to be among them. “Brute forcing” this space to look for anti-cancer activity is not feasible. Even if we had been testing a different compound every second since the universe began, we would still have only tested a minor fraction of this search space. Furthermore, there are immense constraints on how many candidate drugs we can test per unit time. Many chemicals kill cancer cells *in vitro* but the challenge is to find drugs that can kill cancer cells without harming the diversity of normal cells in a living organism. Pre-clinical drug trials, in xenograft or singenic mouse models [135], take time, require very specialist labour and are expensive. And these are just a prerequisite to the further protraction and ginormous expense of clinical trials. Drug development costs are now in the billions of US dollars and these studies take many years [136]. More disturbingly, the drugs – even if found – don’t work well enough: cancer deaths are rising, not falling [1]. Hence, we need exceptional new theory to guide *which* compounds to test. Theoreticians iteratively driving and interpreting experiments. This paper delivers new drug candidates, within a whole new cancer drug class (lipophilic anions), that have an extremely reasoned chance of success. Indeed, the arguments emerge from an experimentally constrained, quantitative framework. Theory has been and is immensely pivotal in physics (i.e. theoretical physics) and its importance is increasingly being realised in biology, in neuroscience especially (e.g. [137-138]), but relatively slowly in cancer biology due to institutional resistance (journals, funding, reviewers etc.). In this paper, and others (pre-published [23, 102] and forthcoming), I have interpreted experiments to produce a new conceptually distinct, explanatory, quantitative, experimentally tractable theory of cancer, which delivers new therapeutic approaches i.e. new, named, chemically-identified drug candidates.

### Caution

The model predicts astronomical therapeutic margins; especially in Figures 2 and 6: thousands, millions, billions, trillions etc. But this is what the Nernst equation, arguably *The* cornerstone of membrane biophysics, predicts with the experimentally derived parameters given to it. If this lipophilic anion approach to cancer fails it will be extremely interesting. It will force the reassessment of the Nernst equation, a fundamental tenant, and/or prompt a review, and perhaps further study, of the experimental values of Ψ_PM_ and Ψ_IM_. This is the distinction between a rational approach, rather than the too-often random approach, to finding new drugs. With rationality: failure can be informative, instructive and guide future direction. Its value then is far beyond just the removal of a molecule out of the immensely, impossibly long list of molecules to trial. Nature and fortuity have gifted us in the past but this low-hanging fruit increasingly looks expended. The rate of drug discovery is slowing, not increasing [139], despite increased R & D spending (Eroom’s law [140]; inverse of Moore’s law).

I am cautiously optimistic that a margin for success may have been hit upon in this work. Even if the Nernst equation loses tenability, the numbers are so large that it only has to fractionally hold to deliver a significant therapeutic margin. Again, if the experimentally recorded Ψ_PM_ and Ψ_IM_ differentials are slightly inaccurate, the accumulation numbers are so large that even if reduced by a substantial fraction: there is still an extant therapeutic margin.

### Conclusion

The present vogue in cancer research, and funding, is to study the different DNA mutations leading to cancer [141]. This study delivers complexity, obfuscation and division because there are many different genetic routes to cancer. Most have interpreted this to mean that there are many different types of cancer. However, I propose that this is incorrect. There is but one end point. The situation is analogous to different neurons, of exactly the same type, employing different ion conductance solutions to reach the same firing pattern, which has been experimentally observed and computationally explored [142]. I suggest that it is better to attack cancer where it arrives, not how it gets there. There are many routes (complexity) but to a single destination (simplicity). As discussed, metastatic cancer ***must*** use a metabolic program distinct from normal adult cells, at least some of the time. This is its point of difference and weakness. The more dangerous the cancer, the greater it’s use of this alternative mode and thence the greater its weakness. The spread of cancer hinges upon its slower ageing/immortality. This is a function of lower ROS generation and greater ROS mitigation. To deny it this, is to beat it. This characteristic must be shared by all metastatic cancers (the relevant, dangerous, perpetuating fraction of it) and is a significant distinction from normal cells. It conveys a cancer-specific Achilles heel. For one thing, as this paper shows, this distinctive program confers cancer cells with a greater accumulation of lipophilic anions than normal cells. This can be leveraged for therapy; and therapy universal to all metastatic cancers.

### Funding

This research was performed unpaid, without funding, and with limited resources in Favela do Terreirão, Rio de Janeiro, Brazil.

## Acknowledgements

This paper is dedicated to the memory of Pte. Conrad Lewis MID, a hero and friend: never forgotten.

## APPENDIX

### A1. Cancer cells may have a depolarised Ψ_OM_ value as compared to normal cells

Victor V. Lemeshko and others have proposed a theory. That Ψ_OM_ = −15 mV in normal cells [54] and Ψ_OM_ = +50 to +60 mV in cancer cells [56-57]. In normal cells, VDAC is open. In cancer cells, VDAC is closed. This closure facilitates aerobic glycolysis in cancer cells. Although, in these cancer cells, a small proportion of their VDAC are opened, in association with the Gibbs free energy of a kinase reaction by hexokinase/glucokinase/creatine kinase [56-57]. And in one version of the model, their binding relates to postulated contact sites between the OM and IM, which bridge the IMS, where an open VDAC in the OM connects with an adenine nucleotide exchanger (ANT) in the IM. More data is needed to assess the voracity of these ideas. The exact Ψ_OM_ value in the Lemeshko cancer cell model [56-57] scales with glucose concentration, tubulin-like effectors and Ψ_IM_ value. When Ψ_IM_ =-160 mV, Ψ_OM_ = ~+55 mV. In this model, the more hyperpolarised Ψ_IM_, the more depolarised Ψ_OM_; so I extrapolate this shown relationship to suggest that even more hyperpolarised values of Ψ_IM_, as is typical of some cancer cells (e.g. Ψ_IM_ =~-210 mV in Neu4145 cancer cells [21]), will scale with even more depolarised values of Ψ_OM_ in cancer cells; further removed from the Ψ_OM_ of normal cells [54]. So, the differential in Ψ_OM_ between normal and cancer cells is large and could become larger still; if the model holds for this wider range. Indeed, this consideration would be an important test of Dr. Lemeshko’s model [56-57]. However, I adhere to his work and, indeed err to underestimate it by not considering any Ψ_OM_ differential between cancer and normal cells to be greater than 50 mV (in either direction). Indeed, the standard version of my model has no Ψ_OM_ differential. My model is independent from, and doesn’t build or rely upon, the Lemeshko model [56-57]. My model is primarily built around Ψ_PM_ and Ψ_IM_ values, which are very experimentally tractable and constrained by data: not Ψ_OM_, which isn’t and is not (to date). But the findings of the Lemeshko model can synergise with the findings of my model when they are incorporated as model parameters (Ψ_OM_ values).

### A2. Why does MKT-077 cause kidney damage?

Kidney cells express *p*-gp. In addition, they express the renal organic cation carrier, which also effluxes cations [79]. Despite this, rhodamine 123 accumulates extensively *ex vivo* in the isolated, perfused rat kidney. 270-360 times the perfusate concentration in kidney tubular cells [79]. Tubular cells uptake the drug passively from the perfusate, driven by their negative-inside plasma membrane potential (-80 mV [79]), and deposit some of it back, actively, to the perfusate. On their opposite face, they deposit it actively into the lumen; to the urine. As the DLC passages across the tubular cell, from the perfusate to urine, it is “actively sequestered” in the tubular cell [79]. This is likely a strategy to maintain a concentration gradient for the DLC into the tubular cell from the perfusate. Inside the tubular cell, the DLC is actively transported into endosomal vesicles in exchange for intravesicular protons [143]. A steep proton gradient is maintained between the vesicle's interior and the cytoplasm by a vesicular proton-ATPase. So, the DLC is not actively imported into the cell directly, but it is indirectly. These kidney cells are distinct from other normal cells in actively importing DLCs. This is perhaps why kidney cells are the first normal cells to be affected by MKT-077. MKT-077 kills cells by causing a general disruption of the mitochondrial inner membrane, undermining the complexes of the respiratory chain within it [34]. MKT-077 conferred membrane disruption may also disrupt these vesicles, and corrupt the activity of their V-ATPase, which will undermine the sequestering of MKT-077 within these kidney cells. Hence MKT-077 can escape from its sequestered shackle and inhibit the respiratory chain. Mortalin is a heat shock protein (Hsp70 family member) that binds and sequesters the p53 transcriptional activator in the cytoplasm. MKT-077 may also harm some normal kidney cells because MKT-077 can prevent this binding and permit an active p53 in the nucleus [33]. Active p53 might switch a cell from aerobic glycolysis to aerobic respiration [144-146]. So, MKT-077 may switch some normal kidney cells out of aerobic glycolysis, which may compromise their hypothesised use of lactate for the urine concentrating mechanism [147]. MKT-077 can’t cross the blood-brain barrier (BBB) [148] and I suggest that it may find future clinical use if it is directly applied to the brain, behind the BBB (e.g. through craniotomy, perhaps after a brain surgery to remove a tumour), where it is insulated from the kidney.

### A3. Cancer cells have a more depolarised Ψ_PM_, perhaps as an adaptation to their greater extracellular acidity: a calculation

Normal cells sit in neutral tissues [119-120]. Cancer cells reside in acidic tumours [119] and have a depolarised Ψ_PM_ relative to normal cells [4-5]. I suggest this is to reduce the proton motive force (pmf) pointing into their cytoplasm, which they must keep neutral like normal cells [47-48, 119-120]. The pmf (Δ*p*) across the plasma membrane has a membrane potential (Δ*ψ*) and pH component (Δ*pH*) [87]. At temperature = 300 K:

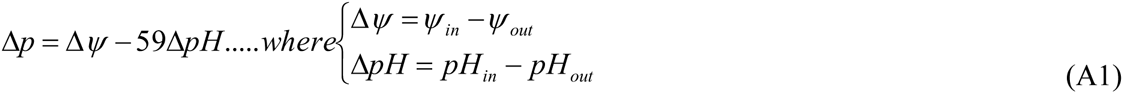

Normal and cancer intracellular pH = 7.2 [47-48, 119-120], extracellular pH to cancer cell = 6.5 [119], extracellular pH to normal cell = 7.4 [119-120], cancer Ψ_PM_ = −10 mV [4], normal Ψ_PM_ = −70 mV [4]:

Normal cell: Δ*p* = 70− (59*(7.4−7.2)) = 58*mV* (Δ*ψ* points inside and Δ*pH* points outside)

Cancer cell: Δp = 10 + (59* (7.2−6.5)) = 51*mV* (Δ*ψ* and Δ*pH* both point inside)

So Δ*p* at the plasma membrane is similar for normal and cancer cells. However, if cancer Ψ_PM_ becomes hyperpolarised to −70 mV, the same as normal cells, Δ*p* more than doubles:

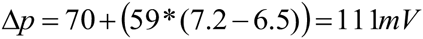

This calculation may help to explain why cancer cells have a depolarised Ψ_PM_ relative to normal cells.

### A4. R code

bulk = 1

valence = 60 # set for a cation; set to −60 for an anion

\# Cancer (Model V1)

CPM = −10

COM = 0

CIM = −200

\# Normal PM = −70

OM = 0

IM = −140

\# NORMAL

cytoplasm = bulk*10^^^(-((((PM-(OM+IM)))/valence)))

intermembrane = bulk*10^^^(-((((PM+OM)-IM)/valence)))

matrix = bulk*10^^^(-((((PM+OM+IM))/valence)))

\# CANCER

C_cytoplasm = bulk*10^^^(-((((CPM-(COM+CIM)))/valence)))

C_intermembrane = bulk*10^^^(-((((CPM+COM)-CIM)/valence)))

C_matrix = bulk*10^^^(-((((CPM+COM+CIM))/valence)))

\# compare

C_cytoplasm/cytoplasm

C_intermembrane/intermembrane

C_matrix/matrix

## REFERENCES

[1] (2012) World Health Organisation (WHO) cancer fact sheet

[2] American Cancer Society. Lifetime Risk of Developing or Dying from Cancer. Available online: http://www.cancer.org/cancer/cancerbasics/lifetime-probability-of-developing-or-dying-from-cancer (accessed on 18 June 2015).

[3] Hille B (2001) Ion channels of excitable membranes. Sunderland, MA: Sinauer.

[4] Yang M, Brackenbury WJ (2013) Membrane potential and cancer progression. Frontiers in physiology 4

[5] Chernet BT, Levin M (2013) Transmembrane voltage potential is an essential cellular parameter for the detection and control of tumor development in a Xenopus model. Disease models & mechanisms 6(3):595–607.

[6] Binggeli R, Cameron IL (1980) Cellular potentials of normal and cancerous fibroblasts and hepatocytes. Cancer research 40(6):1830–1835.

[7] Levin M (2014) Molecular bioelectricity: how endogenous voltage potentials control cell behavior and instruct pattern regulation in vivo. Molecular biology of the cell 25(24):3835–3850.

[8] Levin M (2012) Molecular bioelectricity in developmental biology: new tools and recent discoveries. Bioessays 34(3):205–217.

[9] Pai VP, Martyniuk CJ, Echeverri K, Sundelacruz S, Kaplan DL, Levin M (2015) Genome-wide analysis reveals conserved transcriptional responses downstream of resting potential change in Xenopus embryos, axolotl regeneration, and human mesenchymal cell differentiation. Regeneration.

[10] Sundelacruz S, Levin M, Kaplan DL (2009) Role of membrane potential in the regulation of cell proliferation and differentiation. Stem cell reviews and reports 5(3):231–246.

[11] Bernal SD, Lampidis TJ, Summerhayes IA, Chen LB (1982) Rhodamine-123 selectively reduces clonal growth of carcinoma cells in vitro. Science 218(4577):1117–1119.

[12] Summerhayes IC, Lampidis TJ, Bernal SD, Nadakavukaren JJ, Nadakavukaren KK, Shepherd EL, Chen LB (1982) Unusual retention of rhodamine 123 by mitochondria in muscle and carcinoma cells. Proceedings of the National Academy of Sciences 79(17):5292–5296.

[13] Nadakavukaren KK, Nadakavukaren JJ, Chen LB (1985) Increased rhodamine 123 uptake by carcinoma cells. Cancer research 45(12 Part 1):6093–6099.

[14] Fantin VR, Berardi MJ, Scorrano L, Korsmeyer SJ, Leder P (2002) A novel mitochondriotoxic small molecule that selectively inhibits tumor cell growth. Cancer cell 2(1):29–42.

[15] Modica-Napolitano JS, Aprille JR (2001) Delocalized lipophilic cations selectively target the mitochondria of carcinoma cells. Advanced drug delivery reviews 49(1):63–70.

[16] Davis S, Weiss MJ, Wong JR (1985) Mitochondrial and plasma membrane potentials cause unusual accumulation and retention of rhodamine 123 by human breast adenocarcinoma-derived MCF-7 cells. Journal of Biological Chemistry 260(25): 13844–13850.

[17] Houston MA, Augenlicht LH, Heerdt BG (2011) Stable differences in intrinsic mitochondrial membrane potential of tumor cell subpopulations reflect phenotypic heterogeneity. International journal of cell biology 2011.

[18] Heerdt BG, Houston MA, Augenlicht LH (2005) The intrinsic mitochondrial membrane potential of colonic carcinoma cells is linked to the probability of tumor progression. Cancer Res 65: 9861–9867.

[19] Heerdt BG, Houston MA, Augenlicht LH. Growth properties of colonic tumor cells are a function of the intrinsic mitochondrial membrane potential. Cancer Res 2006;66(3):1591–6.

[20] Chen LB (1988) Mitochondrial membrane potential in living cells. Annual review of cell biology 4(1):155–181.

[21] Fantin VR, St-Pierre J, Leder P (2006) Attenuation of LDH-A expression uncovers a link between glycolysis, mitochondrial physiology, and tumor maintenance. Cancer cell 9(6):425–434.

[22] Bonnet S, Archer SL, Allalunis-Turner J, Haromy A, Beaulieu C et al. (2007) A mitochondria-K+ channel axis is suppressed in cancer and its normalization promotes apoptosis and inhibits cancer growth. Cancer cell 11(1):37–51.

[23] Forrest MD (2015) Why cancer cells have a more hyperpolarised mitochondrial membrane potential and emergent prospects for therapy. bioRxiv 025197. doi: http://dx.doi.org/10.1101/025197

[24] Weiss MJ, Wong JR, Ha CS, Bleday R, Salem RR, Steele GD, Chen LB (1987) Dequalinium, a topical antimicrobial agent, displays anticarcinoma activity based on selective mitochondrial accumulation. Proceedings of the National Academy of Sciences 84(15):5444–5448.

[25] Sun X, Wong JR, Song K, Hu J, Garlid KD, Chen LB (1994) AA1, a newly synthesized monovalent lipophilic cation, expresses potent in vivo antitumor activity. Cancer research 54(6):1465–1471.

[26] Modica-Napolitano JS, Weiss MJ, Chen LB, Aprille JR (1984) Rhodamine 123 inhibits bioenergetic function in isolated rat liver mitochondria. Biochemical and biophysical research communications 118(3):717–723.

[27] Chiba Y, Kubota T, Watanabe M, Matsuzaki SW, Otani Y, Teramoto T,… & Kitajima M (1997) MKT-077, localized lipophilic cation: antitumor activity against human tumor xenografts serially transplanted into nude mice. Anticancer research 18(2A):1047–1052.

[28] Weisberg EL, Koya K, Modica-Napolitano J, Li Y, Chen LB (1996) In vivo administration of MKT-077 causes partial yet reversible impairment of mitochondrial function. Cancer Res 56: 551–555.

[29] Jara JA, Castro-Castillo V, Saavedra-Olavarría, J., Peredo, L., Pavanni, M., Jaña, F.,… & Ferreira, J. (2014). Antiproliferative and uncoupling effects of delocalized, lipophilic, cationic gallic acid derivatives on cancer cell lines. Validation in vivo in singenic mice. Journal of medicinal chemistry 57(6):2440–2454.

[30] Lampidis TJ, Bernal SD, Summerhayes IC, Chen LB (1983) Selective toxicity of rhodamine 123 in carcinoma cells in vitro. Cancer Research 43(2):716–720.

[31] Bernal SD, Lampidis TJ, McIsaac RM, Chen LB (1983) Anticarcinoma activity in vivo of rhodamine 123, a mitochondrial-specific dye. Science 222(4620):169–172.

[32] Herr HW, Huffman JL, Huryk R, Heston WD, Melamed MR, Whitmore WF (1988) Anticarcinoma activity of rhodamine 123 against a murine renal adenocarcinoma. Cancer research 48(8):2061–2063.

[33] Wadhwa R, Sugihara T, Yoshida A, Nomura H, Reddel RR, et al. (2000) Selective toxicity of MKT-077 to cancer cells is mediated by its binding to the hsp70 family protein mot-2 and reactivation of p53 function. Cancer Res 60: 6818–6821.

[34] Modica-Napolitano JS, Koya K, Weisberg E, Brunelli BT, Li Y, Chen LB (1996) Selective damage to carcinoma mitochondria by the rhodacyanine MKT-077. Cancer research 56(3):544–550.

[35] Modica-Napolitano JS, Weissig V (2015) Treatment Strategies that Enhance the Efficacy and Selectivity of Mitochondria-Targeted Anticancer Agents. International journal of molecular sciences 16(8):17394–17421.

[36] Modica-Napolitano JS, Aprille JR (1987) Basis for the selective cytotoxicity of rhodamine 123. Cancer research 47(16):4361–4365.

[37] Wong JR, Ho C, Mauch P, Coleman N, Berman S, Chen LB (1995) Removal of carcinoma cells from contaminated bone marrow using the lipophilic cation rhodamine 123. Clinical cancer research 1(6):621–630.

[38] Dairkee SH, Hackett AJ (1991) Differential retention of rhodamine 123 by breast carcinoma and normal human mammary tissue. Breast cancer research and treatment, 18(1):57–61.

[39] Trapp S, Horobin RW (2005) A predictive model for the selective accumulation of chemicals in tumor cells. European Biophysics Journal 34(7):959–966.

[40] Koya K, Li Y, Wang H, Ukai T, Tatsuta N, Kawakami M, Chen LB (1996) MKT-077, a novel rhodacyanine dye in clinical trials, exhibits anticarcinoma activity in preclinical studies based on selective mitochondrial accumulation. Cancer research 56(3):538–543.

[41] Propper DJ, Braybrooke JP, Taylor DJ, Lodi R, Styles P, Cramer JA,… & Harris AL (1999) Phase I trial of the selective mitochondrial toxin MKT 077 in chemo-resistant solid tumours. Annals of oncology 10(8):923–927.

[42] Britten CD, Rowinsky EK, Baker SD, Weiss GR, Smith L, Stephenson J,… & Eckhardt SG (2000) A phase I and pharmacokinetic study of the mitochondrial-specific rhodacyanine dye analog MKT 077. Clinical cancer research 6(1):42–49.

[43] Jones LW, Narayan KS, Shapiro CA, Sweatman TW (2005) Rhodamine-123: therapy for hormone refractory prostate cancer, a phase I clinical trial. Journal of chemotherapy 17(4):435–440.

[44] Bond SR, Naus CC (2014) The pannexins: past and present. Frontiers in physiology 5.

[45] Bagkos G, Koufopoulos K, Piperi C (2014) A new model for mitochondrial membrane potential production and storage. Medical hypotheses 83(2):175–181.

[46] Bagkos G, Koufopoulos K, Piperi C (2014) ATP Synthesis Revisited: New Avenues for the Management of Mitochondrial Diseases. Current pharmaceutical design 20(28):4570–4579.

[47] Porcelli AM, Ghelli A, Zanna C, Pinton P, Rizzuto R, Rugolo M (2005) pH difference across the outer mitochondrial membrane measured with a green fluorescent protein mutant. Biochemical and biophysical research communications 326(4):799–804.

[48] Davies KM, Strauss M, Daum B, Kief JH, Osiewacz HD, Rycovska A, Kühlbrandt W (2011) Macromolecular organization of ATP synthase and complex I in whole mitochondria. Proceedings of the National Academy of Sciences 108(34):14121–14126.

[49] Cherepanov DA, Feniouk BA, Junge W, Mulkidjanian AY (2003) Low dielectric permittivity of water at the membrane interface: effect on the energy coupling mechanism in biological membranes. Biophysical journal 85(2): 1307–1316.

[50] Llopis J, McCaffery JM, Miyawaki A, Farquhar M, Tsien RY (1998) Measurement of cytosolic, mitochondrial, and Golgi pH in single living cells with green fluorescent proteins, Proc. Natl. Acad. Sci. USA 95: 6803–6808.

[51] Lemeshko SV, Lemeshko VV (2000) Metabolically derived potential on the outer membrane of mitochondria: a computational model. Biophysical journal 79(6):2785–2800.

[52] Lemeshko VV (2002) Model of the outer membrane potential generation by the inner membrane of mitochondria. Biophysical journal 82(2):684–692.

[53] Lemeshko SV, Lemeshko VV (2004) Energy flux modulation on the outer membrane of mitochondria by metabolically-derived potential. Molecular and cellular biochemistry 256(1-2):127–139.

[54] Lemeshko VV (2006) Theoretical evaluation of a possible nature of the outer membrane potential of mitochondria. European Biophysics Journal 36(1):57–66.

[55] Rostovtseva T, Colombini M (1997) VDAC channels mediate and gate the flow of ATP: implications for the regulation of mitochondrial function. Biophysical journal 72(5):1954.

[56] Lemeshko VV (2014) VDAC electronics: 1. VDAC-hexo (gluco) kinase generator of the mitochondrial outer membrane potential. Biochimica et Biophysica Acta (BBA)-Biomembranes 1838(5): 1362–1371.

[57] Lemeshko VV (2014) VDAC electronics: 2. A new, anaerobic mechanism of generation of the membrane potentials in mitochondria. Biochimica et Biophysica Acta (BBA)-Biomembranes 1838(7): 1801–1808.

[58] Lipinski CA, Lombardo F, Dominy BW, Feeney PJ (2012) Experimental and computational approaches to estimate solubility and permeability in drug discovery and development settings. Advanced drug delivery reviews 64: 4–17.

[59] National Center for Biotechnology Information. PubChem Compound Database; CID=60099122, https://pubchem.ncbi.nlm.nih.gov/compound/60099122 (accessed Dec 18, 2015)

[60] National Center for Biotechnology Information. PubChem Compound Database; CID=73357197, https://pubchem.ncbi.nlm.nih.gov/compound/73357197 (accessed Dec 18, 2015)

[61] Tabet S, Douglas SF, Daze KD, Garnett GA, Allen KJ, Abrioux EM,… & Hof F (2013) Synthetic trimethyllysine receptors that bind histone 3, trimethyllysine 27 (H3K27me3) and disrupt its interaction with the epigenetic reader protein CBX7. Bioorganic & medicinal chemistry 21(22):7004–7010.

[62] National Center for Biotechnology Information. PubChem Compound Database; CID=45271389, https://pubchem.ncbi.nlm.nih.gov/compound/45271389 (accessed Dec 18, 2015)

[63] Inglese J, Auld DS, Jadhav A, Johnson RL, Simeonov A, Yasgar A,… & Austin CP (2006) Quantitative high-throughput screening: a titration-based approach that efficiently identifies biological activities in large chemical libraries. Proceedings of the National Academy of Sciences 103(31):11473–11478.

[64] R Core Team (2013) R: A language and environment for statistical computing. R Foundation for Statistical Computing, Vienna, Austria. URL: http://www.R-project.org

[65] Zamzami N, Kroemer G (2001) The mitochondrion in apoptosis: how Pandora’s box opens. Nat. Rev. Mol. Cell Biol. 2: 67–71

[66] Kim S, Thiessen PA, Bolton EE, Chen J, Fu G, Gindulyte A,… & Bryant SH (2015) PubChem Substance and Compound databases. Nucleic Acids Research:gkv951.

[67] Saengkhae C, Loetchutinat C, Garnier-Suillerot A (2003) Kinetic analysis of rhodamines efflux mediated by the multidrug resistance protein (MRP1).Biophysical journal 85(3):2006–2014.

[68] Kurtoglu M, Lampidis TJ (2009) From delocalized lipophilic cations to hypoxia: blocking tumor cell mitochondrial function leads to therapeutic gain with glycolytic inhibitors. Molecular nutrition & food research 53(1):68–75.

[69] Lizard G, Chignol MC, Chardonnet Y, Schmitt D (1994) Active cell membrane mechanisms involved in the exclusion of RH 123 allow distinction between normal and tumoral cells. Cell biology and toxicology 10(5–6):399–406.

[70] Pusztai L, Wagner P, Ibrahim N, Rivera E, Theriault R, Booser D,… & Hortobagyi, GN (2005) Phase II study of tariquidar, a selective p-glycoprotein inhibitor, in patients with chemotherapy-resistant, advanced breast carcinoma. Cancer 104(4):682–691.

[71] Thomas H, Coley HM (2003) Overcoming multidrug resistance in cancer: an update on the clinical strategy of inhibiting p-glycoprotein. Cancer control 10(2): 159–159.

[72] Shukla S, Wu CP, Ambudkar SV (2008) Development of inhibitors of ATP-binding cassette drug transporters-present status and challenges.

[73] Brouty-Boye D, Kolonia D, Wu CJ, Savaraj N, Lampidis TJ. Relationship of multidrug resistance to rhodamine-123 selectivity between carcinoma and epithelial cells: Taxol and vinblastine modulate drug efflux. Cancer Res. 1995;55:1633–1638.

[74] Kramer R, Weber TK, Morse B, Arceci R, Staniunas R, Steele Jr G, Summerhayes IC (1993) Constitutive expression of multidrug resistance in human colorectal tumours and cell lines. British journal of cancer, 67(5), 959.

[75] Cordon-Cardo C, O'brien JP, Boccia J, Casals D, Bertino JR, Melamed MR (1990) Expression of the multidrug resistance gene product (p-glycoprotein) in human normal and tumor tissues. Journal of Histochemistry & Cytochemistry 38(9):1277–1287.

[76] Fojo AT, Ueda KSDJ, Slamon DJ, Poplack DG, Gottesman MM, Pastan I (1987) Expression of a multidrug-resistance gene in human tumors and tissues. Proceedings of the National Academy of Sciences 84(1):265–269.

[77] David-Beabes GL, Overman MJ, Petrofski JA, Campbell PA, de Marzo AM, Nelson WG (2000) Doxorubicin-resistant variants of human prostate cancer cell lines DU 145, PC-3, PPC-1, and TSU-PR1: characterization of biochemical determinants of antineoplastic drug sensitivity. International journal of oncology 17(6): 1077–1163.

[78] Tatsuta N, Suzuki N, Mochizuki T, Koya K, Kawakami M, Shishido T,… & Chen LB (1999) Pharmacokinetic analysis and antitumor efficacy of MKT-077, a novel antitumor agent. Cancer chemotherapy and pharmacology 43(4):295–301.

[79] Masereeuw R, Moons MM, Russel FG (1997) Rhodamine 123 accumulates extensively in the isolated perfused rat kidney and is secreted by the organic cation system. European journal of pharmacology 321(3):315–323.

[80] Gamboa-Vujicic G, Emma DA, Liao SY, Fuchtner C, Manetta A (1993) Toxicity of the mitochondrial poison dequalinium chloride in a murine model system. Journal of pharmaceutical sciences 82(3):231–235.

[81] Kruh GD, Belinsky MG (2003) The MRP family of drug efflux pumps. Oncogene 22(47):7537–7552.

[82] Leslie EM, Deeley RG, Cole SP (2005) Multidrug resistance proteins: role of P-glycoprotein, MRP1, MRP2, and BCRP (ABCG2) in tissue defense. Toxicology and applied pharmacology 204(3):216–237.

[83] Inui KI, Masuda S, Saito H (2000) Cellular and molecular aspects of drug transport in the kidney. Kidney international 58(3):944–958.

[84] Balendiran GK, Dabur R, Fraser D (2004) The role of glutathione in cancer. Cell biochemistry and function 22(6):343–352.

[85] Stryer L, Berg JM, Tymoczko JL (2002) Biochemistry, 4^th^ Ed. New York, NY: WH Freeman.

[86] Alberts B, Johnson A, Lewis J, Raff M, Roberts K, Walter P (1994) Molecular Biology Of The Cell, 3^rd^ Ed. New York, NY: Garland Publishing.

[87] Nicholls DG, Ferguson S (2013) Bioenergetics. Academic Press.

[88] Warburg O (1956) On the origin of cancer cells. Science 123(3191):309–314.

[89] Ferreira LM (2010) Cancer metabolism: the Warburg effect today. Exp Mol Pathol 89(3):372–380.

[90] Kim JW, Dang CV (2006) Cancer’s molecular sweet tooth and the Warburg effect. Cancer Res 66(18):8927–8930.

[91] Lopez-Lazaro M (2008) The warburg effect: why and how do cancer cells activate glycolysis in the presence of oxygen? Anti-Cancer Agent Me 8(3):305–312.

[92] Elliott RL, Jiang XP, Head JF (2014) Want to Cure Cancer? Then Revisit the Past; “Warburg Was Correct”, Cancer Is a Metabolic Disease. Journal of Cancer Therapy

[93] Gatenby RA, Gillies RJ (2004) Why do cancers have high aerobic glycolysis? Nat Rev Cancer 4(11):891–899.

[94] Bui T, Thompson CB (2006) Cancer’s sweet tooth. Cancer cell 9(6):419–420.

[95] Nakashima RA, Paggi MG, Pedersen PL (1984) Contributions of glycolysis and oxidative phosphorylation to adenosine 5'-triphosphate production in AS-30D hepatoma cells. Cancer Res 44(12 Part 1):5702–5706.

[96] Diaz-Ruiz R, Rigoulet M, Devin A (2011) The Warburg and Crabtree effects: On the origin of cancer cell energy metabolism and of yeast glucose repression. BBA-Bioenergetics 1807(6):568–576.

[97] DeBerardinis RJ, Lum JJ, Hatzivassiliou G, Thompson CB (2008) The biology of cancer: metabolic reprogramming fuels cell growth and proliferation. Cell metabolism, 7(1):11–20.

[98] Mathupala SP, Ko YH, Pedersen PL (2010) The pivotal roles of mitochondria in cancer: Warburg and beyond and encouraging prospects for effective therapies. Biochimica et Biophysica Acta (BBA)-Bioenergetics 1797(6):1225–1230.

[99] Vander Heiden MG, Cantley LC, Thompson CB (2009) Understanding the Warburg effect: the metabolic requirements of cell proliferation. Science 324(5930):1029–1033.

[100] Czernin J, Phelps ME (2002) Positron emission tomography scanning: current and future applications. Annu Rev Med 53: 89–112.

[101] Gambhir SS, Czernin J, Schwimmer J, Silverman DH, Coleman RE, Phelps MEA (2001) Tabulated summary of the FDG PET literature. J Nucl Med 42:1S–93S.

[102] Forrest MD (2015) NADH as a cancer medicine. bioRxiv:019307. doi: http://dx.doi.org/10.1101/019307

[103] Birsoy K, Possemato R, Lorbeer FK, Bayraktar EC, Thiru P et al. (2014) Metabolic determinants of cancer cell sensitivity to glucose limitation and biguanides. Nature 508(7494):108–112.

[104] Moreno-Sánchez R, Rodríguez-Enríquez S, Saavedra E, Marín-Hernández A, Gallardo-Pérez JC (2009) The bioenergetics of cancer: is glycolysis the main ATP supplier in all tumor cells? Biofactors 35(2): 209–225.

[105] Wadhwa R, Sugihara T, Yoshida A, Nomura H, Reddel RR, et al. (2000) Selective toxicity of MKT-077 to cancer cells is mediated by its binding to the hsp70 family protein mot-2 and reactivation of p53 function. Cancer Res 60: 6818–6821.

[106] Schulz TJ, Thierbach R, Voigt A, et al (2006) Induction of oxidative metabolism by mitochondrial frataxin inhibits cancer growth: Otto Warburg revisited. J Biol Chem 281: 977–81.

[107] Christofk HR, Vander Heiden MG, Harris MH, Ramanathan A, Gerszten RE, Wei R,… & Cantley LC (2008) The M2 splice isoform of pyruvate kinase is important for cancer metabolism and tumour growth. Nature 452(7184):230–233.

[108] Alves S (2008) Two of a kind. Nature Reports Stem Cells.

[109] Baker M (2008) Cancer and embryonic stem cells share genetic fingerprints. Nature Reports Stem Cells.

[110] Chung S, Dzeja PP, Faustino RS, Perez-Terzic C, Behfar A, Terzic A (2007) Mitochondrial oxidative metabolism is required for the cardiac differentiation of stem cells. Nat Clin Pract Cardiovasc Med 4: S60—S67.

[111] Teslaa T, Teitell MA (2014) Pluripotent stem cell energy metabolism: an update. The EMBO journal e201490446.

[112] Zalzman M, Falco G, Sharova LV, Nishiyama A, Thomas M, Lee SL,… & Ko MS (2010) Zscan4 regulates telomere elongation and genomic stability in ES cells. Nature 464(7290):858–863.

[113] Hayflick L (1997) Mortality and immortality at the cellular level. A review. Biochemistry-New York-English Translation of Biokhimiya 62(11):1180–1190.

[114] Kondoh H, Lleonart ME, Bernard D, Gil J (2007) Protection from oxidative stress by enhanced glycolysis; a possible mechanism of cellular immortalization. Histology and histopathology 22(1): 85–90.

[115] Hong Y, Stambrook PJ (2004) Restoration of an absent G1 arrest and protection from apoptosis in embryonic stem cells after ionizing radiation. PNAS 101(40):14443–14448.

[116] De Grey AD (1999) The mitochondrial free radical theory of aging. Austin, TX: RG Landes.

[117] Blacker TS, Mann ZF, Gale JE, Ziegler M, Bain AJ, Szabadkai G, Duchen MR (2014) Separating NADH and NADPH fluorescence in live cells and tissues using FLIM. Nature communications 5.

[118] Yu Q, Heikal AA (2009) Two-photon autofluorescence dynamics imaging reveals sensitivity of intracellular NADH concentration and conformation to cell physiology at the single-cell level. Journal of Photochemistry and Photobiology B: Biology 95(1):46–57.

[119] Damaghi M, Wojtkowiak JW, Gillies RJ (2013) pH sensing and regulation in cancer. Frontiers in physiology 4.

[120] Casey JR, Grinstein S, Orlowski J (2010) Sensors and regulators of intracellular pH. Nature reviews Molecular cell biology 11(1):50–61.

[121] Brooks GA (2010) What does glycolysis make and why is it important? Journal of applied physiology 108(6):1450–1451.

[122] McCarty MF, Whitaker J (2010) Manipulating tumor acidification as a cancer treatment strategy. Altern Med Rev 15(3):264–72.

[123] Dejan Zecevic et al. (2009) Voltage-sensitive dye. Scholarpedia 4(3):3355.

[124] Walton RD, Mitrea BG, Pertsov AM, Bernus O (2009) A novel near-infrared voltage-sensitive dye reveals the action potential wavefront orientation at increased depths of cardiac tissue. In Engineering in Medicine and Biology Society, 2009. EMBC 2009. Annual International Conference of the IEEE (pp. 4523–4526). IEEE.

[125] Bagkos G, Koufopoulos K, Piperi C (2015) Mitochondrial emitted electromagnetic signals mediate retrograde signaling. Medical hypotheses 85(6):810–818.

[126] Atkins P, De Paula J (2011) Physical chemistry for the life sciences. Oxford University Press.

[127] Takahara K, Inamoto T, Minami K, Yoshikawa Y, Takai T, Ibuki N, et al. (2015) The Anti-Proliferative Effect of Boron Neutron Capture Therapy in a Prostate Cancer Xenograft Model. PLoS ONE 10(9):e0136981.

[128] Dewi N, Yanagie H, Zhu H, Demachi K, Shinohara A, Yokoyama K,… & Takahashi H (2013) Tumor growth suppression by gadolinium-neutron capture therapy using gadolinium-entrapped liposome as gadolinium delivery agent. Biomedicine & Pharmacotherapy 67(6):451–457.

[129] Leford H (2015) World’s largest cancer charity lays out field’s grand challenges. Nature 526: 301–302

[130] Pathak RK, Marrache S, Harn DA, Dhar S (2014) Mito-DCA: a mitochondria targeted molecular scaffold for efficacious delivery of metabolic modulator dichloroacetate. ACS chemical biology 9(5):1178–1187.

[131] Debele TA, Peng S, Tsai HC (2015) Drug Carrier for Photodynamic Cancer Therapy. International journal of molecular sciences 16(9):22094–22136.

[132] National Center for Biotechnology Information. PubChem Compound Database; CID=4254851, https://pubchem.ncbi.nlm.nih.gov/compound/4254851 (accessed Dec 17, 2015)

[133] Apostolova N, Victor VM (2015) Molecular Strategies for Targeting Antioxidants to Mitochondria: Therapeutic Implications. Antioxidants & Redox Signaling. 22(8):686–729.

[134] Virshup AM, Contreras-García J, Wipf P, Yang W, Beratan DN (2013) Stochastic voyages into uncharted chemical space produce a representative library of all possible drug-like compounds. Journal of the American Chemical Society 135(19):7296–7303.

[135] Huang C-Y, Huang H-Y, Forrest MD, Pan Y-R, Wu W-J, Chen H-M (2014) Inhibition Effect of a Custom Peptide on Lung Tumors. PLoS ONE 9(10):e109174.

[136] Mullard A (2014) New drugs cost US$2.6 billion to develop. Nature Reviews Drug Discovery 13: 877

[137] Forrest MD (2015) Simulation of alcohol action upon a detailed Purkinje neuron model and a simpler surrogate model that runs> 400 times faster. BMC neuroscience 16(1):27.

[138] Forrest MD (2014) Intracellular calcium dynamics permit a Purkinje neuron model to perform toggle and gain computations upon its inputs. Frontiers in computational neuroscience 8.

[139] Pammolli F, Magazzini L, Riccaboni M (2011) The productivity crisis in pharmaceutical R&D. Nature reviews Drug discovery 10(6):428–438.

[140] Scannell JW, Blanckley A, Boldon H, Warrington B (2012) Diagnosing the decline in pharmaceutical R&D efficiency. Nature reviews Drug discovery 77(3):191–200.

[141] Wagstaff A (2013) Jim Watson: DNA revealed the causes, it may never reveal a cure. CancerWorld. September-October.

[142] Achard P, De Schutter E (2006) Complex parameter landscape for a complex neuron model. PLoS Comput Biol 2(7):e94.

[143] Pritchard JB, Miller DS (1997) Renal secretion of organic cations: a multistep process. Advanced drug delivery reviews 25(2):231–242.

[144] Matoba S, Kang JG, Patino WD, Wragg A, Boehm M, Gavrilova O,… & Hwang, P. M (2006) p53 regulates mitochondrial respiration. Science 312(5780):1650–1653.

[145] Ma W, Sung HJ, Park JY, Matoba S, Hwang PM (2007) A pivotal role for p53: balancing aerobic respiration and glycolysis. Journal of bioenergetics and biomembranes 39(3):243–246.

[146] Wang L, Schumann U, Liu Y, Prokopchuk O, Steinacker JM (2012) Heat shock protein 70 (Hsp70) inhibits oxidative phosphorylation and compensates ATP balance through enhanced glycolytic activity. Journal of Applied Physiology 113(11):1669–1676.

[147] Hervy S, Thomas SR (2003) Inner medullary lactate production and urine-concentrating mechanism: a flat medullary model. American Journal of Physiology-Renal Physiology 284(1):F65–F81.

[148] Miyata Y, Li X, Lee HF, Jinwal UK, Srinivasan SR, Seguin SP, Gestwicki JE (2013) Synthesis and initial evaluation of YM-08, a blood-brain barrier permeable derivative of the heat shock protein 70 (Hsp70) inhibitor MKT-077, which reduces tau levels. ACS chemical neuroscience 4(6):930–939.

